# The ELF3 transcription factor is associated with an epithelial phenotype and represses epithelial-mesenchymal transition

**DOI:** 10.1101/2022.08.19.504435

**Authors:** Ayalur Raghu Subbalakshmi, Sarthak Sahoo, Prakruthi Manjunatha, Shaurya Goyal, Vignesh A Kasiviswanathan, M Yeshwanth, R Soundharya, Isabelle McMullen, Jason A. Somarelli, Mohit Kumar Jolly

**Affiliations:** Centre for BioSystems Science and Engineering, Indian Institute of Science, Bangalore 560012, India; Department of Medical Electronics, M S Ramaiah Institute of Technology, Bangalore 560054, India; Department of Humanities and Social Sciences, Faculty of Sciences, Indian Institute of Technology, Kharagpur 721302, India; Department of Biotechnology, JSS Science and Technology University, Mysore 570006, India; Department of Biotechnology, National Institute of Technology Warangal, 506004, India; Department of Medicine, Duke University, Durham, NC 27708, USA; Duke Cancer Institute, Duke University, Durham, NC 27708, USA

**Author notes:** Authors to whom correspondence should be addressed (J.A.S), (M.K.J). These authors contributed equally (Ayalur Raghu Subbalakshmi, Sarthak Sahoo).

**Keywords:** ELF3, phenotypic plasticity, mathematical modeling, epithelial-mesenchymal transition (EMT), Mesenchymal–Epithelial Transition (MET)

## Abstract

Epithelial-mesenchymal plasticity (EMP) involves bidirectional transitions between epithelial, mesenchymal and multiple intermediary hybrid epithelial/mesenchymal phenotypes. While the process of epithelial-mesenchymal transition (EMT) and its associated transcription factors are well-characterised, the transcription factors that promote mesenchymal-epithelial transition (MET) and stabilise hybrid E/M phenotypes are less well understood. Here, we analyse multiple publicly-available transcriptomic datasets at bulk and single-cell level and pinpoint ELF3 as a factor that is strongly associated with an epithelial phenotype and is inhibited during EMT. Using mechanism-based mathematical modelling, we also show that ELF3 inhibits the progression of EMT, suggesting ELF3 may be able to counteract EMT induction, including in the presence of EMT-inducing factors, such as WT1. Our model predicts that the MET induction capacity of ELF3 is stronger than that of KLF4, but weaker than that of GRHL2. Finally, we show that ELF3 levels correlates with worse patient survival in a subset of solid tumor types, suggesting cell-of-origin or lineage specificity in the prognostic capacity of ELF3.

## Introduction

Phenotypic plasticity – the ability of cancer cells to reversibly change their phenotypes to adapt to changing environments – is crucial for cancer cell survival. It is a hallmark of metastasizing cancer cells that enables them to alter their cell-cell adhesion and migration traits, evade the immune system, and resist targeted therapies (Celià-Terrassa and Kang, 2016; Gupta *et al*., 2019). Given the importance of phenotypic plasticity as a critical regulator of metastasis and therapy resistance, there is a crucial need to decode the dynamics of phenotypic plasticity in cancer.

Epithelial-mesenchymal transition (EMT) and its reverse - mesenchymal-epithelial transition (MET) – constitute a key axis of phenotypic plasticity, through bidirectional transitions between epithelial, mesenchymal, and one or more hybrid epithelial/mesenchymal (E/M) phenotype(s) (Pastushenko and Blanpain, 2019; Tripathi *et al*., 2020). Once tacitly assumed to be a binary process, now EMT is conceptualized as a spectrum of cell states, with many manifestations of the highly plastic and heterogeneous hybrid E/M phenotypes (Pastushenko *et al*., 2018; Cook and Vanderhyden, 2020; Lourenco *et al*., 2020; Deshmukh *et al*., 2021). Many EMT-inducing transcription factors (EMT-TFs), such as ZEB1/2, SNAI1/2, and TWIST have been well-characterised (Peinado *et al*., 2007; Taube *et al*., 2010; Drápela *et al*., 2020), but TFs that can stabilize hybrid E/M phenotypes or induce MET are less well characterized. Most of the MET-TFs identified to date – e.g. GRHL1/2, OVOL1/2 and KLF4 – induce MET by forming mutually inhibitory feedback loops with EMT-TFs (Xiang *et al*., 2012, 2017; Roca *et al*., 2013; Somarelli *et al*., 2016; Fujimoto *et al*., 2019; Watanabe *et al*., 2019; Yang *et al*., 2019; Subbalakshmi *et al*., 2022a). Similarly, while time-course transcriptomic bulk and single-cell data on EMT has been now extensively collected, the dynamics of MET remains less well-studied (Zhang *et al*., 2014; Celià-Terrassa *et al*., 2018; Karacosta *et al*., 2019; Stylianou *et al*., 2019; Cook and Vanderhyden, 2020). Given the proposed roles of MET in metastatic colonization and therapeutic response, a better understanding of MET and its regulators is needed.

Among the potential candidate transcription factors that may promote MET, the transcription factor E74-like factor 3 (ELF3) belongs to the E26 transformation-specific (ETS) family of transcription factors. It is strongly expressed in epithelial tissues, such as the digestive tract, bladder, and lungs, where it plays key roles in differentiation and homeostasis (Suzuki *et al*., 2021). It has also been shown to inhibit EMT in multiple cancer types. For instance, in bladder cancer cells, overexpression of ELF3 reduced invasion and expression of mesenchymal markers (Gondkar *et al*., 2019). Similarly, ELF3 correlated with an epithelial phenotype in ovarian cancer cells, and its overexpression in SKOV3 cells reduced invasion and led to a downregulation of mesenchymal markers and an increase in epithelial markers (Yeung *et al*., 2017), reminiscent of observations made in lung cancer cells (Lou *et al*., 2018). In colorectal cancer, knockdown of ELF3 in HCT116 cells induced ZEB1 upregulation. ELF3 expression was found to antagonize ZEB1 expression by inhibiting the Wnt and RAS oncogenic signalling pathways (Liu *et al*., 2019). Consistent reports in non-transformed mouse mammary gland epithelial cell line (NMuMG) showed that ELF3 correlated strongly with E-cadherin (*Cdh1*) expression and led to activation of *Grhl3* (Sengez *et al*., 2019), thereby playing an important role as gatekeeper of an epithelial lineage. Together, these studies suggest that ELF3 may be a putative MET-TF.

At a molecular level, ELF3 is inhibited by both the SNAI family members SNAI1 (SNAIL) and SNAI2 (SLUG) (Lyons *et al*., 2008; Li *et al*., 2021a), both of which can induce EMT to varying degrees (Bolós *et al*., 2003; Subbalakshmi *et al*., 2022b). ELF3, in turn, can repress upregulation of ZEB1/2 by ETS1 in breast cancer (Sinh *et al*., 2017), head and neck squamous carcinoma (Sakamoto *et al*., 2021) and in normal bile duct epithelial cells (Suzuki *et al*., 2021). ESE1 and ETS1 are dominantly present in luminal and basal-like subtypes of breast cancer cells, and reciprocally regulate each other, thus impacting the EMT status of these cells (Sinh *et al*., 2017). Moreover, similar to ZEB1 (Jolly *et al*., 2018), ELF3 can self-activate (Li *et al*., 2021b).

Here, we utilize the experimental observations discussed above, along with multiple transcriptomic data sets to develop a mechanism-based mathematical model to delineate the impact of ELF3 on epithelial-mesenchymal plasticity. Our model predicts that ELF3 can delay or prevent the onset of EMT; consequently, its overexpression can induce a partial or complete MET. Analysis of publicly-available *in vitro* transcriptomics data, including that from the Cancer Cell Line Encyclopedia (CCLE), and The Cancer Genome Atlas (TCGA) revealed that ELF3 is negatively correlated with mesenchymal factors and positively correlated with epithelial factors. Further, analysis of time-course transcriptomic data shows that ELF3 levels decrease upon EMT induction, which further supports the hypothesis that ELF3 acts as a putative MET-TF. Finally, ELF3 levels are associated with cancer patient survival in a lineage- and cancer-specific manner, highlighting the clinical relevance of ELF3 in specific cancer types.

## Results

### ELF3 is associated with an epithelial phenotype

We first investigated the association between ELF3 expression levels and both epithelial and mesenchymal programs across cancer cell lines. In the CCLE cohort, we quantified the correlation coefficient for each individual gene with epithelial and mesenchymal scores using single-sample gene expression enrichment (ssGSEA) (Tan *et al*., 2014) (**Fig 1A**). As expected, the mesenchymal genes VIM, ZEB1, SNAI1 and SNAI2 were positively correlated with mesenchymal ssGSEA scores and negatively correlated with epithelial scores. Conversely, the canonical epithelial genes CDH1, GRHL2 and OVOL2 showed a strong positive correlation with ssGSEA-based epithelial scores and negative correlation with ssGSEA-based mesenchymal scores. ELF3 was present among the epithelial factors (**Fig 1A**), reminiscent of its previously-reported positive correlation with *Cdh1* and negative correlation with *Vim* (Sengez *et al*., 2019; Watanabe *et al*., 2019). Next, we examined the correlation of ELF3 with these scores in the CCLE cohort in a cancer type-specific manner (**Fig 1B**). We observed that in a majority of cancer types, including breast cancer, prostate cancer and bladder cancer, ELF3 correlated positively with epithelial scores and negatively with mesenchymal scores. These trends were consistent in TCGA cancer types as well (**Fig 1C**), further suggesting that ELF3 correlates with an epithelial phenotype.

**Figure 1:**
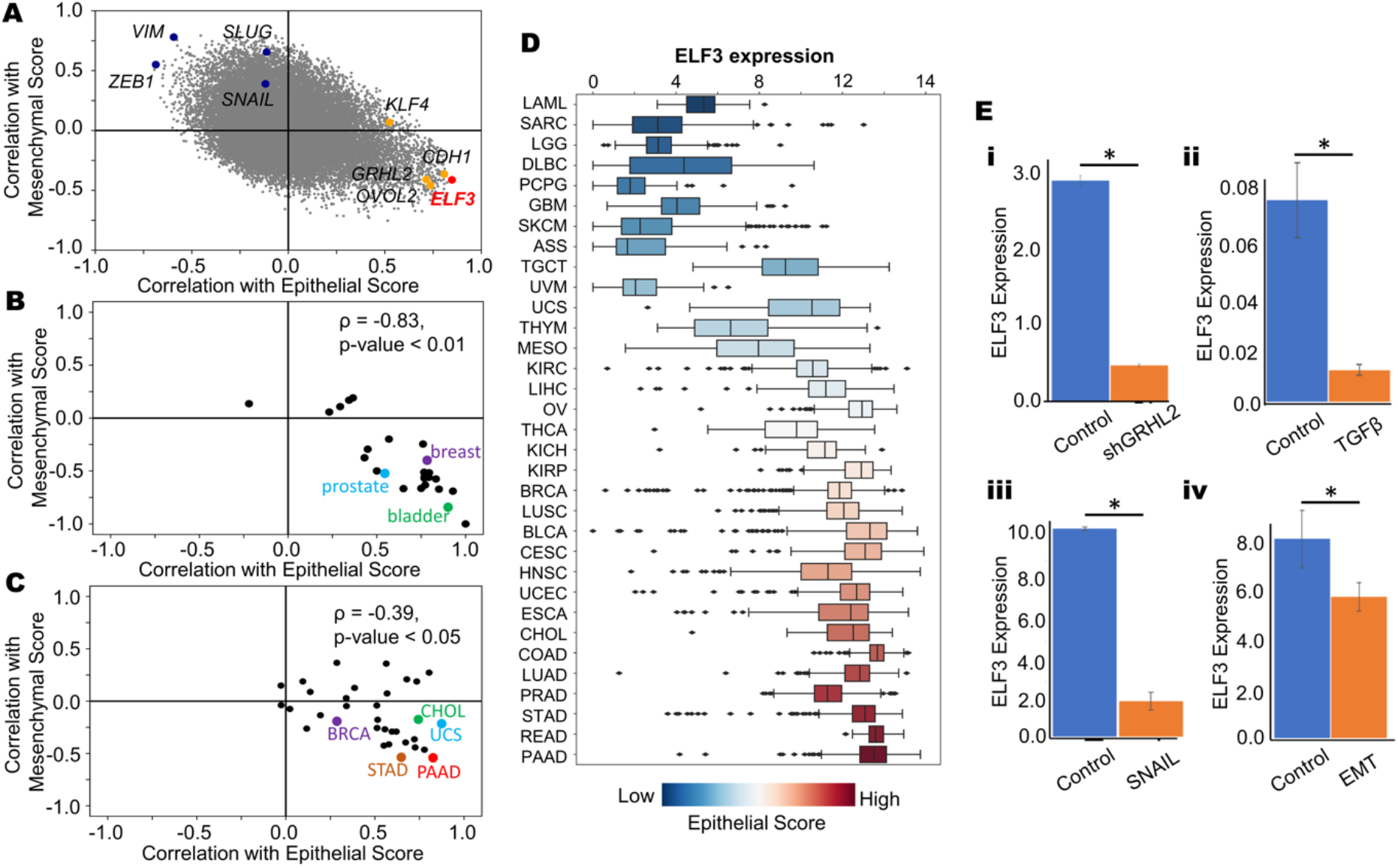
ELF3 correlates with an epithelial phenotype. **A)** Scatterplot showing the correlation coefficients of individual genes with epithelial and mesenchymal scores across the CCLE cohort. Mesenchymal genes VIM, ZEB1, SNAI1 and SNAI2 are represented in blue and epithelial genes GRHL2, OVOL2, KLF4 and CDH1 are represented in orange. ELF3 is represented in red. **B)** Tissue specific correlations of ELF3 with epithelial and mesenchymal scores in the CCLE cohort when grouped by tissue of origin. **C)** Correlations of ELF3 with epithelial and mesenchymal scores across different TCGA cancer types. **D)** Boxplot showing ELF3 expression levels across different cancer types in TCGA. Cancer types are ordered by increasing median epithelial scores. **E)** Changes in ELF3 expression during EMT and/or MET induction across GEO datasets. I) GSE118407 ii) GSE61220 iii) GSE58252 iv) GSE59922. *: p< 0.05 (Students’ t-test).

We next tabulated ELF3 expression levels with respect to the median epithelial ssGSEA scores in a given cancer type. We observed that an increase in ELF3 expression levels was concordant with that in the corresponding median epithelial scores (**Fig 1D**). Conversely, a decrease in ELF3 levels coincided with increase in EMT scores (**Fig S1A**), thereby highlighting that ELF3 expression levels are higher in epithelial cancer types (PAAD: pancreatic adenocarcinoma, STAD: stomach adenocarcinoma, READ: rectum adenocarcinoma, PRAD: prostate adenocarcinoma, LUAD: lung adenocarcinoma) when compared to mesenchymally-derived cancer types (SARC: sarcoma, LGG: low grade glioma, GBM: glioblastoma) (**Fig 1D**). We next compared the methylation status of ELF3 in comparison to TCGA samples. We observed that the methylation status of ELF3 correlated negatively with its expression, and the methylation was usually higher in mesenchymal cancer types (**Fig S1B**). Together, these analyses suggest that ELF3 strongly correlates with an epithelial state across cancers.

Next, we asked whether ELF3 levels are downregulated during EMT, using publicly-available transcriptomics datasets. We first examined changes in ELF3 expression levels in response to silencing of GRHL2 in OVCA4209 cells (GSE118407) which led to induction of EMT (Chung *et al*., 2019) and reduction in ELF3 levels (**Fig 1E, i**). Similarly, in TGFβ-induced EMT in airway epithelial cells (Tian *et al*., 2015) ELF3 levels were downregulated (**Fig 1E, ii**; GSE61220). Consistent trends were observed in MCF-7 cells that were forced to undergo EMT by the overexpression of SNAIL (McGrail *et al*., 2015) (GSE58252; **Fig 1E, iii**), and in mouse mammary EpRas cells undergoing a TGFβ-driven EMT (GSE59922; **Fig 1E, iv**) (Johansson *et al*., 2015). Together, these observations indicate that downregulation of ELF3 is a consistent marker of EMT.

### ELF3 is inhibited during EMT induction and can prevent EMT

We next investigated temporal changes in ELF3 expression levels in time-course transcriptomic datasets. A549 lung adenocarcinoma cells treated with TGFβ to undergo EMT (GSE17708; **Fig 2A**) (Sartor *et al*., 2009) showed a progressive decrease in ELF3 levels at later time-points of induction. ELF3 expression was also strongly negatively correlated with the enrichment of the Hallmark EMT signature (r = -0.91, p < 0.001). Next, we interrogated ELF3 levels in LNCaP prostate cancer cells along the EMT trajectory upon SNAIL induction and a subsequent MET over 20 days after withdrawal of SNAIL induction (Stylianou *et al*., 2019). ELF3 levels were reduced during EMT progression and re-expressed during MET induction (GSE80042; **Fig 2B**). SNAIL- and TGFβ-induced EMT in MCF10A breast epithelial (Comaills *et al*., 2016) also led to reduction in ELF3, irrespective of the mode of EMT induction (GSE89152**; Fig 2C**). We also analysed ELF3 expression in single-cell RNA-seq data in samples treated with TGFβ for a period of seven days to undergo EMT followed by three days of recovery for cells to undergo MET (Cook and Vanderhyden, 2020). Across multiple cell lines – A549 (left), DU145 (center) and OVCA420 (right) – ELF3 expression levels are inhibited with the onset of EMT, but a recovery in ELF3 expression is observed as they undergo MET (GSE147405, **Fig 2D**). Together, these analyses suggest that ELF3 is inhibited in a reversible manner during induction of EMT across multiple contexts.

**Figure 2:**
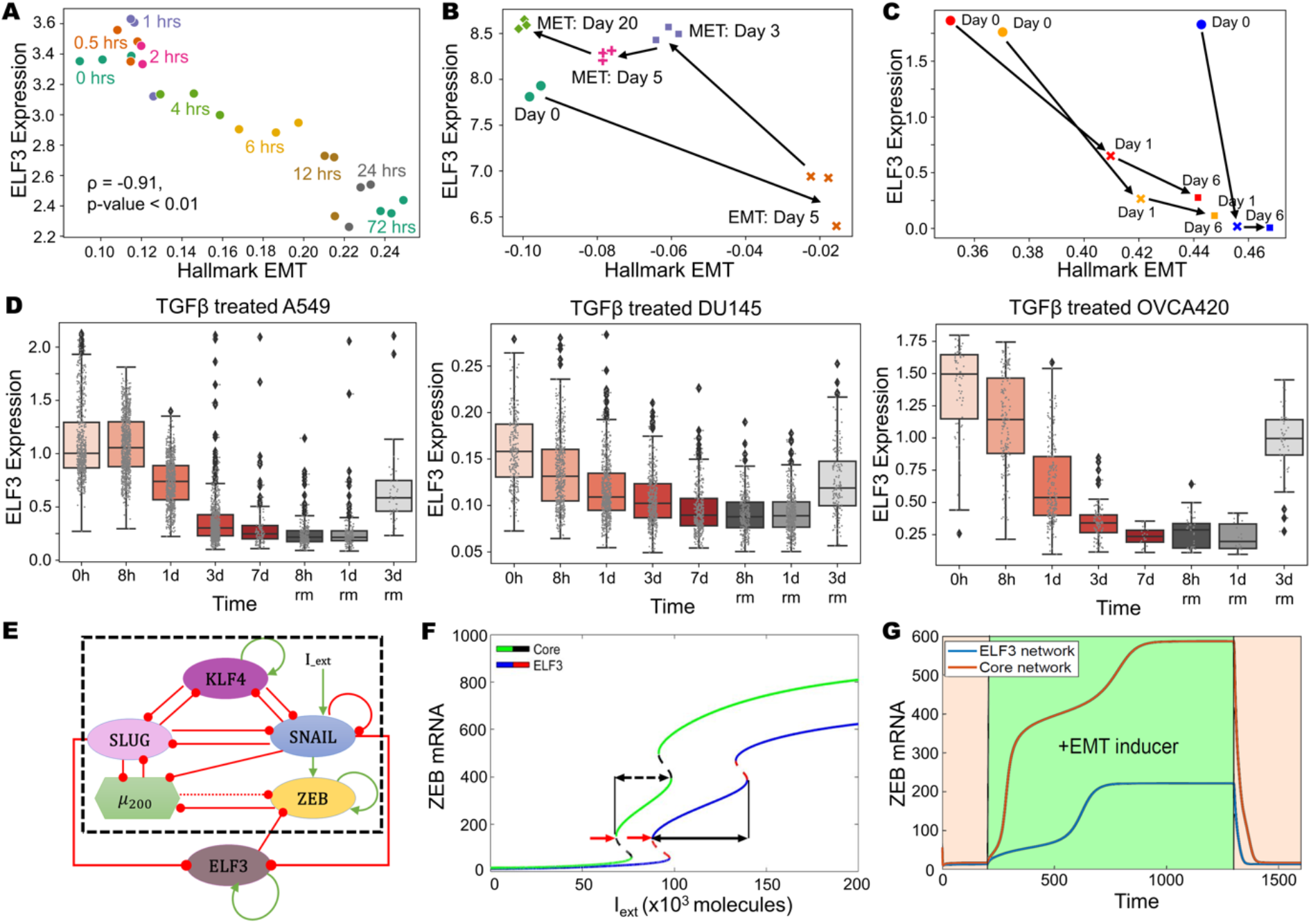
Analysis of ELF3 levels during induction of EMT and/or MET. **A)** Scatterplot of ssGSEA scores for the “Hallmark EMT” pathway with ELF3 expression levels at different time points of EMT induction (GSE17708). Spearman’s correlation coefficient and corresponding p-value is given. **B)** Scatterplot and trajectory of samples in terms of ssGSEA scores of Hallmark EMT with ELF3 expression in EMT induction via SNAIL over expression (5 days) and subsequent induction of MET over a 20-day period (GSE80042). **C)** Same as B) but for treatment with TGFβ (red, orange profiles) and SNAIL induction (blue profile) over a 6-day time period (GSE89152). **D)** Single-cell data of ELF3 levels in TGFβ treated A549 (left), DU145 (center) and OVCA420 (right) over a 7 day period (EMT) followed by TGFβ withdrawal (MET) for the next 3 days (GSE147405). **E)** Schematic representation of ELF3 coupled with an EMT regulatory network (dotted rectangle) consisting of miR-200, ZEB1, SNAIL, SLUG and KLF4. Green arrows denote activation, and red bars indicate inhibition. Solid arrows represent transcriptional regulation; dotted lines represent microRNA-mediated regulation. **F**) Bifurcation diagrams for ZEB1/2 mRNA levels in response to an external signal (I_ext) levels for the coupled EMT–ELF3 circuit (solid blue and dotted red curve) and the core EMT circuit (solid green and dotted black curve). Black arrows indicate the region of the hybrid E/M state and red arrows indicate a switch from an epithelial phenotype. **G)** Temporal dynamics of ZEB1/2 mRNA levels in a cell starting in an epithelial phenotype when exposed to a high level of an external EMT signal (I_ext = 100,000 molecules) (green-shaded region) for the circuits shown in panel E.

Next, we examined the role of ELF3 in modulating EMT dynamics. We analyzed the interaction dynamics between ELF3 and a core EMT regulatory circuit (denoted by black dotted rectangle in **Fig 2E**) comprised of five core factors: three EMT-inducing transcription factors (EMT-TFs) - ZEB1/2, SNAIL, and SLUG - and two EMT-inhibiting factors: the microRNA miR-200 family (Gregory *et al*., 2008) and KLF4, a transcription factor that correlates with the epithelial phenotype (Yori *et al*., 2010; Subbalakshmi *et al*., 2021). First, we plotted a bifurcation diagram to track the levels of ZEB1/2 mRNA (as a readout of EMT phenotype) in response to an external EMT-inducing signal I_ext (**Fig 2F**). With an increase in I_ext levels, cells switched from an epithelial state (low levels of ZEB1/2 mRNA) to a hybrid E/M phenotype (moderate levels of ZEB1/2 mRNA) and, finally, to a mesenchymal state (high levels of ZEB1/2 mRNA). In the absence of ELF3 (curve with green solid line and black dashed line), the switch from an epithelial to mesenchymal phenotype occurred at a much lower strength of I_ext than when compared to the network that contained ELF3 (curve with blue solid line and red dashed line) (indicated using red arrows) (**Fig 2F**). In addition, in the presence of ELF3, the region of I_ext for which the hybrid E/M state existed was larger when compared to the core network (dotted black arrows), indicating that ELF3 can stabilize a hybrid E/M state.

We further mapped the temporal response for a fixed value of I_ext signal. We noted a transition from an epithelial state first to a hybrid E/M state and then to a mesenchymal state in response to I_ext. However, in the presence of ELF3, this transition was more gradual and relatively slower as compared to the absence of ELF3 (blue curve vs. red curve in **Fig 2G**). Consistently, the steady-state value of ZEB1/2 mRNA levels seen in the presence of ELF3 was relatively lower, due to ELF3-mediated inhibition of ZEB1/2. This trend can also be corroborated by reduced ZEB1/2 levels in the bifurcation diagram (blue curve lies below green curve at all values of I_ext in **Fig 2F**).

We next estimated the extent to which ELF3 impacted EMT dynamics depending on the strength of its interactions with the EMT circuit. When the strength of repression of ZEB1/2 mRNA by ELF3 was reduced, we observed an expansion of the {M} region (a mesenchymal phenotype) accompanied by a shrinking of the {E} (only epithelial) and {H} (only hybrid E/M) regions (**Fig S2A**). Conversely, when the strength of ELF3 self-activation was increased or the repression of SLUG on ELF3 was decreased, it resulted in expansion of the {E} and {H} regions and a reduction of the {M} region (**Fig S2B-C**). No major qualitative changes were observed in network dynamics in the above-mentioned cases. To further evaluate the impact of other kinetic parameters on our model predictions, we performed sensitivity analysis by varying the numerical values of the input kinetic parameters by ±10% one by one and captured the changes in the range of the I_ext values for the existence of the hybrid E/M state in the bifurcation diagram. Except for a few parameters, most of which did not influence the interactions of ELF3 with the core EMT circuit (except threshold value of ZEB1/2 repression), this change did not extend beyond 5-10% (**Fig S2D**). Importantly, an approximately 35% percent decrease in the region of hybrid E/M phenotypes was estimated when ELF3 was not considered in the network. Overall, this analysis indicates that the behaviour of ELF3 in its ability to delay or prevent EMT induction is robust to small parametric variations.

### ELF3 is predicted to act as an MET inducer

To further determine the role of ELF3 in EMT dynamics, we expanded the network to incorporate GRHL2, a potent MET-TF that forms a mutually inhibitory loop with ZEB1 and can activate ELF3 (Chung *et al*., 2016; Farris *et al*., 2016; Jolly *et al*., 2016; Mooney *et al*., 2017) (**Fig S3A**). We simulated the dynamics of this network across an ensemble of parameter values and initial conditions, through RACIPE (Huang *et al*., 2018) and collated all the steady states obtained. In this ensemble of steady states, both ELF3 values and EMT scores (= ZEB1 – miR-200) showed a bimodal distribution (**Fig S3B**). Principal Component Analysis (PCA) reveals two clusters along the PC1 (which explains 53.41% variance), one of which has low EMT scores and high ELF3, while the other has high EMT scores and low ELF3 levels (**Fig 3A, i-ii**). These results suggest that across the parameter sets considered (each of which can be thought of as representing an individual cell in a heterogeneous population), this network can recapitulate E-M heterogeneity. Projecting SLUG levels on the PCA plot revealed that SLUG expression was higher in mesenchymal and hybrid E/M phenotypes (**Fig 3A, iii**). This trend is in concordance with earlier experimental observations that associate SLUG with varying degrees of EMT (Wels *et al*., 2011; Sterneck *et al*., 2020; Subbalakshmi *et al*., 2022b). Finally, we projected the levels of GRHL2, miR-200, ZEB1 and KLF4 individually on the PCA plot. While GRHL2 expression largely mimicked that of miR-200 or ELF3, ZEB1 expression resembled that of an EMT score (**Fig S3C**). However, KLF4 patterns did not completely overlap with other epithelial factors, GRHL2 and ELF3; KLF4 was also high in hybrid E/M phenotypes. This difference indicates a stronger concordance between GRHL2 and ELF3 in associating with an epithelial state (**Fig 3A, iv**). A similar difference was also observed in the CCLE cohort scatter plots for the correlation of individual genes with epithelial and mesenchymal scores, where GRHL2 and ELF3 behaved similarly as potential inhibitors of EMT, but KLF4 did not show any significant association with mesenchymal score (**Fig 1A**).

**Figure 3:**
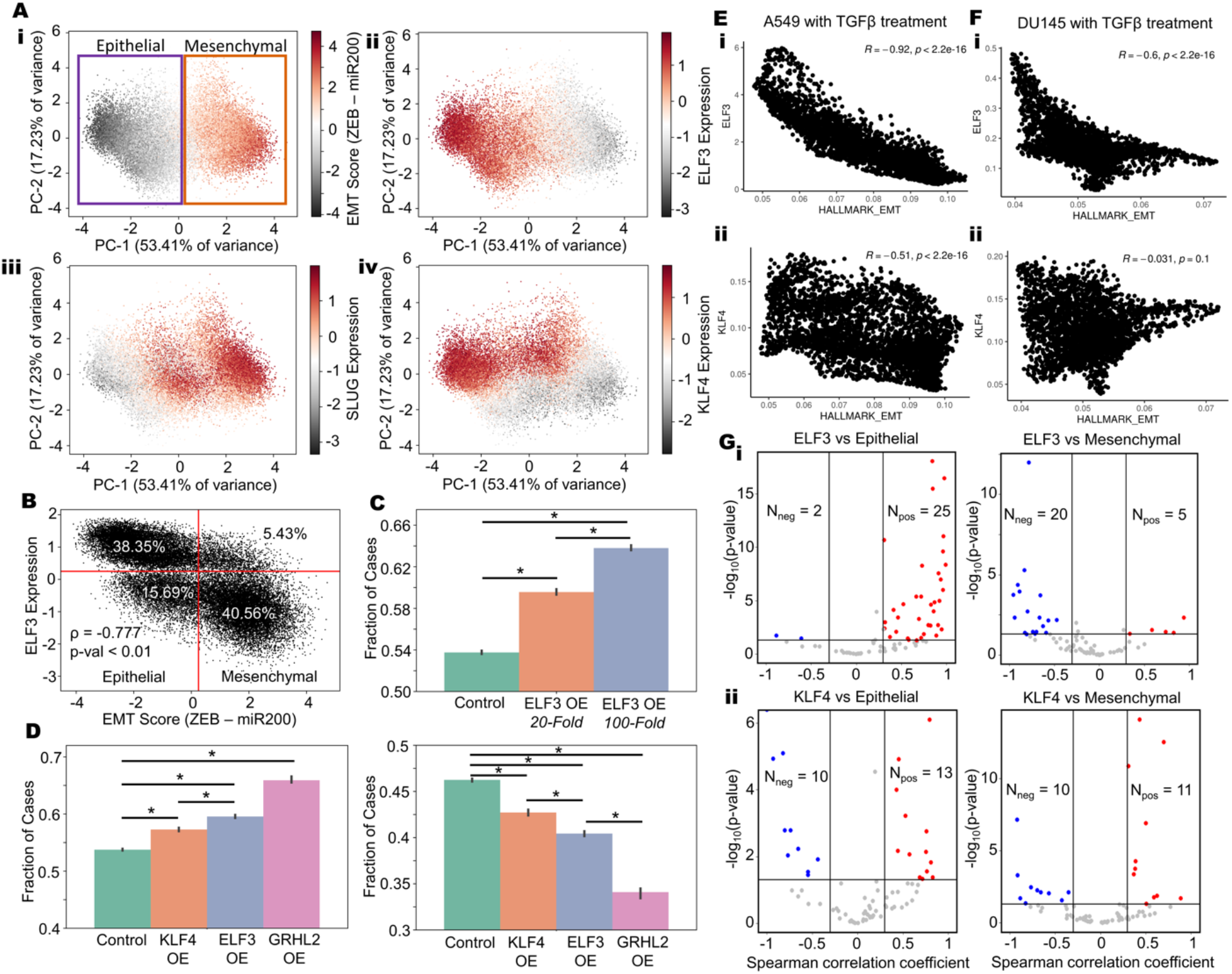
ELF3 as a MET inducer. **A)** PCA scatter plot of all steady states of RACIPE colored by (i) EMT score (= ZEB – miR200), ii) SLUG levels iii) ELF3 levels and iv) KLF4 levels. **B)** Scatterplot of EMT scores and ELF3 levels across steady state solutions obtained from RACIPE. Red lines indicate the position of minima in the bimodal distributions of EMT scores and ELF3 levels. Spearman correlation coefficient and p-value are mentioned. **C)** Fraction of steady state solutions resulting in Epithelial phenotype in control, 20-fold and 100-fold over expression of ELF3. * represents a statistically significant difference in the fraction of cases in the epithelial phenotype (Students’ t-test; p < 0.05). **D)** Fraction of steady state solutions resulting in the Epithelial (left panel) and Mesenchymal (right panel) phenotypes in control, 20-fold over expression of ELF3, GRHL2 and KLF4. *represents a statistically significant difference (Students’ t-test; p < 0.05). **E)** Correlation of ELF3 and KLF4 with Hallmark EMT ssGSEA scores for single-cell RNA-seq data of A549 cells treated with TGFβ. Spearman’s’ correlation coefficient values are mentioned (GSE147405). **F)** Same as E) but for DU145. **G)** Volcano plots showing correlation of ELF3 and KLF4 levels with ssGSEA epithelial and mesenchymal scores in a meta-analysis of breast cancer datasets. Each dot represents a dataset. R < - 0.3, p< 0.05 or R > 0.3, p < 0.05 are counted as statistically significant cases. N_neg_ denotes number of datasets for which a negative correlation (blue dots) is observed, N_pos_ denotes number of datasets for which a positive correlation (red dots) is observed between the two corresponding expression levels or ssGSEA scores.

Based on these observations, we used the bimodally-distributed and inversely-correlated EMT scores and ELF3 expression levels to quantify the *in silico* population distribution of epithelial and mesenchymal phenotypes. For the network shown here, approximately 54% of cells can be classified as epithelial while 46% cells can be binned as mesenchymal (**Fig 3B**). Amongst this population, approximately 71% of the epithelial cells had high levels of ELF3 while only 12% of mesenchymal cells were high in ELF3 expression. This clearly demonstrates that high ELF3 expression is predominantly associated with an epithelial phenotype. Next, we determined the effect of ELF3 overexpression on the system by simulations where we overexpressed ELF3 by 20-fold or by 100-fold. These results showed a dose-dependent and statistically reliable increase in the proportion of cells exhibiting an epithelial state (**Fig 3C**), supporting the notion that ELF3 is an MET inducer. We next compared the MET-inducing capabilities of ELF3 with that of GRHL2 and KLF4 (**Fig 3D**). GRHL2 overexpression resulted in the highest epithelial fraction and the lowest mesenchymal fraction. Following GRHL2, ELF3 was found to be the next most potent inducer, followed by KLF4 as the weakest MET inducer.

To further interrogate this trend, we compared the correlation of ELF3 and KLF4 scores with epithelial (= miR-200 + GRHL2) and mesenchymal (= ZEB + SNAIL + SLUG) factors individually, based on our simulation data. Again, ELF3 showed stronger correlations as compared to KLF4 (**Fig S3D-E**). These *in silico* trends were also recapitulated in single-cell RNA-seq data for A549 and DU145 with TGFβ treatment (Cook and Vanderhyden, 2020) where ELF3 shows stronger trends compared to KLF4 in terms of its correlation with “Hallmark EMT” scores (A549: r = - 0.92 for ELF3 vs. r= - 0.51 for KLF4; DU145: r = - 0.6 for ELF3 vs. r = - 0.03 for KLF4) and with 76-gene signature (76GS)-based scoring of EMT in which higher values indicate an epithelial behavior (Chakraborty *et al*., 2020) (A549: r = 0.92 for ELF3 vs. r = 0.45 for KLF4; DU145: r = 0.48 for ELF3 vs. r = 0.21 for KLF4) (**Fig 3E-F, S4A-B**). Finally, in a meta-analysis across multiple transcriptomic datasets belonging to breast cancer, ovarian cancer and bladder cancer (**Table S1**), we investigated the correlation of ELF3, GRHL2 and KLF4 with epithelial and mesenchymal gene sets. Among the 27 datasets in breast cancer where ELF3 correlated significantly (p < 0.05, r > 0.3 or r < - 0.3) with the epithelial signature, the correlation was positive in 25 datasets. Conversely, among 25 breast cancer datasets where ELF3 correlated significantly with the mesenchymal signature, the correlation was negative in 20 datasets (**Fig 3G**). While GRHL2 showed similar trends as to ELF3, KLF4, on the other hand, did not show such strong trends, across the three cancer types investigated here (**Fig 3G, S4C-D**). Together, these results propose ELF3 as a putative MET-inducer, albeit with potentially weaker MET-inducing capacity than GRHL2.

### Correlation of ELF3 with patient survival

The role of ELF3 as a regulator of epithelial plasticity led us to query whether ELF3 is associated with clinical outcomes in cancer. To do this, we analyzed a series of gene expression data sets across solid tumors. In breast cancer, high ELF3 levels correlated with worse patient outcomes in terms of overall survival, relapse-free survival and metastasis-free survival (**Fig 4A, S5A-C**) (GSE3494, GSE9893, GSE4922, GSE65308 and GSE48408), reminiscent of observations that ELF3 can act as an independent prognostic marker for poor survival in hormone receptor positive (ERα+, PR+) HER2+ breast cancer patients (Kar and Gutierrez-Hartmann, 2017; Kar *et al*., 2020). Similar trends have been observed in prostate cancer (Longoni *et al*., 2013) and non-small cell lung cancer (Wang *et al*., 2018a). However, the trend was reversed in colorectal cancer, where high ELF3 levels correlated with better patient prognosis in terms of overall survival, relapse-free survival and metastasis-free survival (GSE16125, GSE39582, GSE28814 and GSE28722) (**Fig 4B, S5D-F**), similar to reports in ovarian (Yeung *et al*., 2017) and bladder cancer (Gondkar *et al*., 2019). Thus, ELF3 appears to associate with patient survival in a cancer-specific manner.

**Fig 4:**
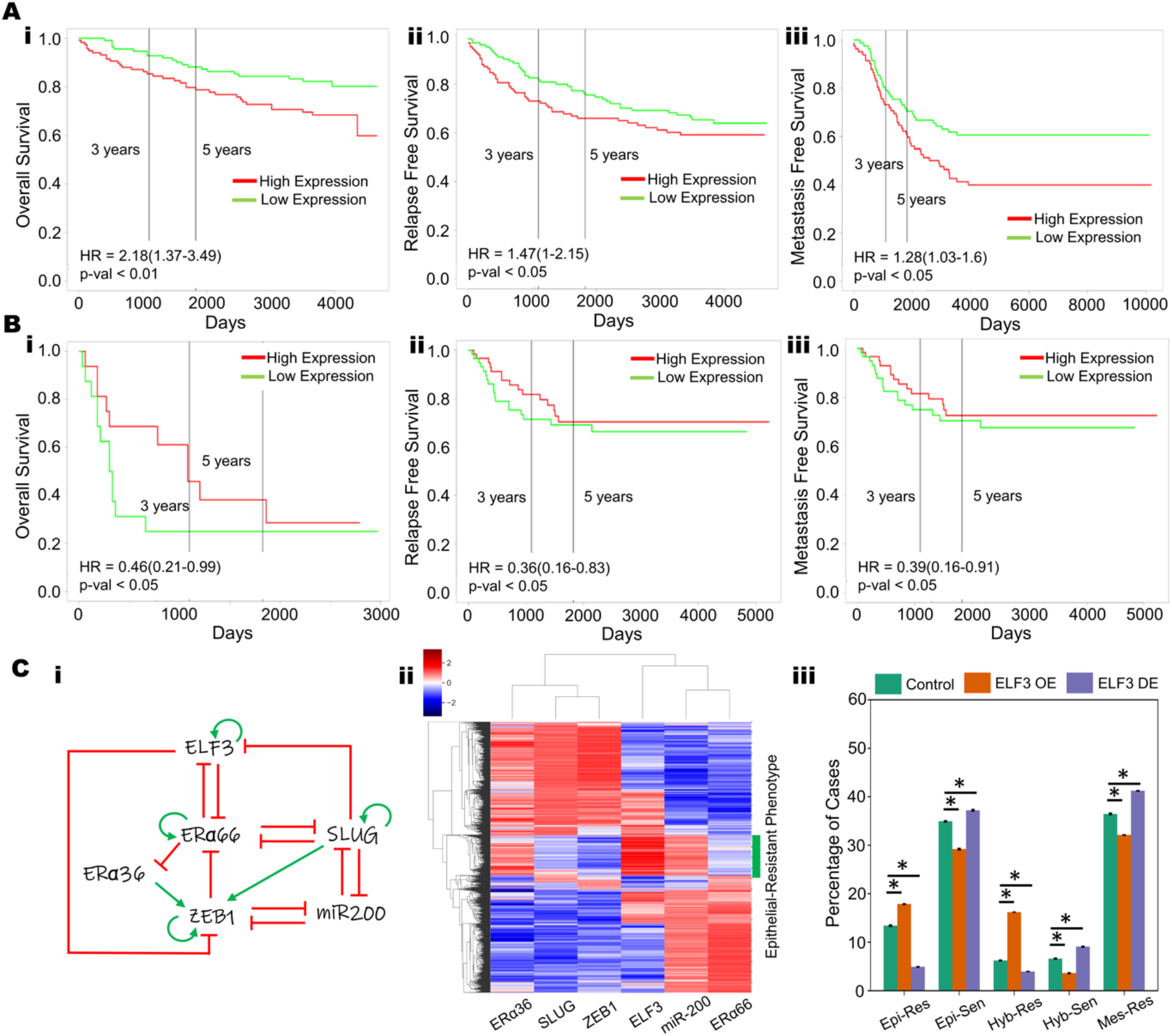
Cancer type-specific correlation of ELF3 with patient survival. **A)** Higher ELF3 levels correlate with worse patient outcomes in breast cancer samples (i) overall survival (GSE3494), ii) relapse-free survival (GSE4922), and iii) metastasis-free survival (GSE48408) **B)** In colorectal cancer samples, ELF3 levels correlate with worse patient outcomes: i) overall survival (GSE16125), ii) relapse-free survival (GSE28814), and iii) metastasis-free survival (GSE28814), showing lower. **C)** i) A gene regulatory network coupling ELF3 with the EMT core network (miR-200, ZEB1, SLUG) and Estrogen Receptor isoforms (ERα66, ERα36) in the context of ER+ breast cancer. Red hammers represent inhibitory links and green arrows represent activation links; ii) Heatmap of steady state solutions upon simulation of the GRN in i); iii) Percentage of steady state solutions resulting in each of the phenotype pairs: Epithelial and Resistant (Epi-Res), Epithelial and Sensitive (Epi-Sen), Hybrid and Resistant (Hyb-Res), Hybrid and Sensitive (Hyb-Sen), and Mesenchymal and Resistant (Mes-Res) in control, 20-fold up or downregulation of ELF3; * represents a statistically significant difference in the fraction of cases that end up in the epithelial phenotype (Students t-test; p-val < 0.05).

To gain further insights into this context-specific behavior, we focused on ER+ (estrogen receptor positive) breast cancer. Earlier work, including ours, has shown that in ER+ breast cancer, EMT and tamoxifen resistance can promote each other (Hiscox *et al*., 2006; Tian and Schiemann, 2017; Wang *et al*., 2019; Sahoo *et al*., 2021). With this in mind, we investigated how ELF3 may influence the EMT-tamoxifen resistance interaction. Our mechanism-based model for coupling EMT factors (miR-200, ZEB, SLUG) with two isoforms of ER (ERα66 and ERα36) had predicted that while the predominant phenotypes are either epithelial/tamoxifen-sensitive or mesenchymal/tamoxifen-resistant, there are also other states that can be observed, including epithelial/tamoxifen-resistant, hybrid (E/M)/tamoxifen-resistant and hybrid(E/M)/tamoxifen-sensitive (Sahoo *et al*., 2021). Thus, we incorporated experimentally-identified connections of ELF3 with ERα66 and ERα36 into our coupled EMT-ELF3 network and simulated the dynamics of this ER+ breast cancer-specific network using RACIPE (Huang *et al*., 2018). In the ER+ breast cancer context, ELF3 can repress the transcriptional function of ERα66 (Gajulapalli *et al*., 2016), similar to the role of its family member ELF5, which can suppress ERα66 and its downstream targets, thus mediating tamoxifen resistance in luminal breast cancer cells (Kalyuga *et al*., 2012). Conversely, ELF3 is known to be inhibited by ERα66 in MCF7 and ZR-75.1 cells (Cicatiello *et al*., 2010), thereby potentially forming a mutually inhibitory loop (**Fig 4C**, i).

Simulation of this gene regulatory network (**Fig 4C**, ii) using RACIPE suggests that it can enable an epithelial-like, tamoxifen-sensitive state characterized by high levels of miR-200 and ERα66; low levels of SLUG, ZEB1 and ERα36 and a mesenchymal-like, tamoxifen-resistant state characterized by low levels of miR-200 and ERα66; high levels of SLUG, ZEB1 and ERα36. We also observed that a subset of the epithelial cluster, with high expression of miR-200 and ZEB1 is associated with high expression of ELF3, which is consistent with the role of ELF3 in promoting an epithelial-like phenotype. However, this cluster had a significantly lower expression of ERα66 and a higher expression of ERα36 (**Fig 4C**, ii). As ERα66 is the target of anti-estrogen drugs, such as tamoxifen, the loss or downregulation of ERα66 is often associated with a more resistant phenotype. Conversely, upregulation ERα36 is associated with a tamoxifen-resistant phenotype (Wang *et al*., 2018b). The association of ELF3 with this epithelial phenotype that also exhibits a more resistant phenotype may be one of the key contributing factors that explain the relationship between ELF3 and worse survival in breast cancer. To further substantiate the role of ELF3, we mimicked ELF3 overexpression *in silico* and found that it increased the frequency of an epithelial/ tamoxifen-resistant phenotype comprised of high levels of miR-200 and ERα36 and low levels of ZEB1 and ERα66, while that of epithelial/tamoxifen-sensitive phenotype decreased. Conversely, downregulating ELF3 showed opposite trends (**Fig 4C**, iii). While additional experimental data supporting this hypothesis is needed to validate the importance of these relationships in tamoxifen resistance, the observed upregulation of another ETS family member, ELF5, in tamoxifen-resistant MCF7 cells (Kalyuga *et al*., 2012; Fitzgerald *et al*., 2016) and tamoxifen-resistant brain metastases (Piggin *et al*., 2020), as well as differential expression of ELF3 in tamoxifen-treated vs. control groups (Gielen *et al*., 2005), lends credence to this hypothesis.

### ELF3 can inhibit EMT that is mediated by factors such as WT1

Given the proposed role of ELF3 in safeguarding an epithelial phenotype, we analysed whether ELF3 can prevent EMT induction when an additional factor is added to the abovementioned regulatory network. As an example of an additional EMT inducing factor, we focused on Wilms Tumour (WT1). WT1 was found to transcriptionally repress *Cdh1* and activate *Snail* in epicardial cells, where its knockdown reduced the frequency of cardio-vascular progenitor cells and its derivatives (Martínez-Estrada *et al*., 2010). Similarly, in NSCLC (non-small cell lung cancer) and prostate cancer, WT1 inhibits *Cdh1* and promotes invasion (Brett *et al*., 2013; Wu *et al*., 2013). WT1 levels were found to be higher in cancer cells relative to cancer-adjacent, non-tumor tissue, while CDH1 levels were lower in the cancer cells as compared to the cancer-adjacent tissue (Wu *et al*., 2013; Han *et al*., 2020). Similarly, in breast cancer, WT1-positive tumors were found to be more mesenchymal, and overexpression of WT1 in breast epithelial cells, HBL100, led to upregulation of mesenchymal markers, such as Vimentin (*Vim*) and Tenascin C (*Tnc*) (Artibani *et al*., 2017). Together, these observations highlight WT1 as a potent EMT-inducer.

At a molecular level, WT1 is self-inhibitory (Reddy *et al*., 1995), while promoting the expression of SNAIL (Martínez-Estrada *et al*., 2010) and inhibiting the expression of SLUG (Takeichi *et al*., 2013). Based on these experimental data, we expanded our network model to incorporate these interactions (**Fig 5A**). Next, we calculated the bifurcation diagram of ZEB1/2 mRNA levels in response to an external EMT-inducing signal I_ext, for four different circuits: core network (no ELF3, no WT1: WT1-/ELF3-), core network + ELF3 (WT-/ELF3+), core network + ELF3 + WT1 (WT1+/ELF3+), core network + WT1 (WT+/ELF3-) (**Fig 5B**). The first two bifurcation diagrams (WT-/ELF3-, WT-/ELF3+ – shown in green solid and black dotted curve, and blue solid and red dotted curve respectively) are the same as we calculated earlier (**Fig 2D**), showing that the presence of ELF3 required more I_ext to force cells out of an epithelial phenotype. In scenarios of WT1-/ELF3+ (blue solid and red dashed curve), and WT1-/ELF3- (solid green and black dashed curve), the switch from an epithelial to mesenchymal phenotype occurred at a much higher strength of I_ext than when compared to the network which contained WT1, either in presence (solid purple and black dashed curve) or absence (solid yellow with red dashed curve) of ELF3 (WT1+/ELF3-, WT1+/ELF3+) (**Fig 6B**). Further, in the presence of WT1 (indicated by solid black arrow), the region of I_ext for which the hybrid E/M state existed shrunk when compared to the network containing ELF3 but not WT1 (indicated by dotted black arrow), further indicating that ELF3 can inhibit WT1-induced EMT.

**Figure 5:**
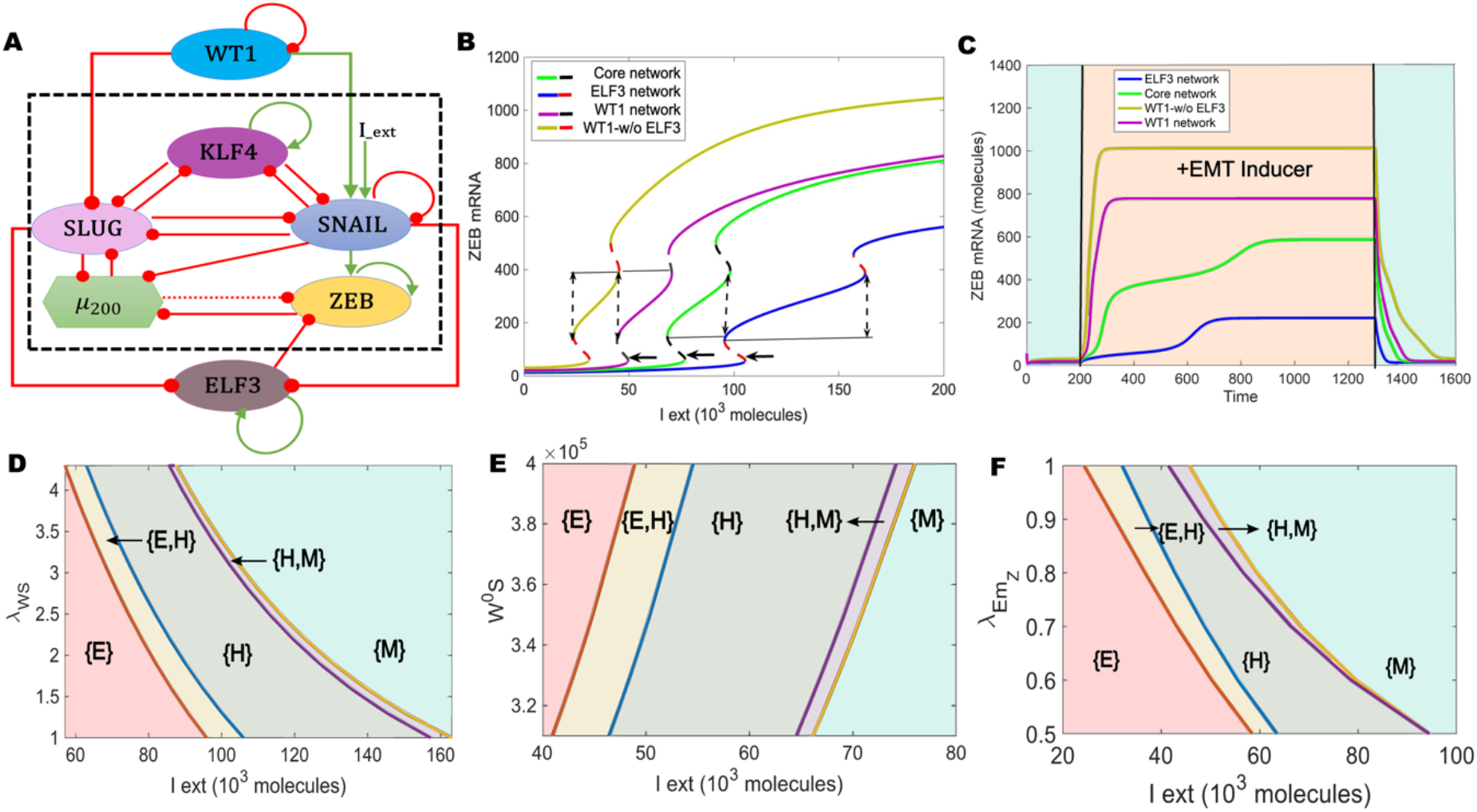
ELF3 can inhibit induction of EMT by WT1. **A)** Schematic representation of the ELF3 network coupled with WT1. Green arrows denote activation, red bars indicate inhibition. **B)** Bifurcation diagrams for ZEB1/2 mRNA levels in response to an external signal (I_ext) levels for the coupled WT1 coupled ELF3 network (solid pink and dotted black curve), WT1 coupled with the core EMT circuit (no ELF3) (solid yellow and dotted red curve), EMT–ELF3 circuit (solid blue and dotted red curve) and core EMT circuit (solid green and dotted black curve). Solid lines indicate the region of the hybrid state; arrows indicate the switch from epithelial phenotype. **C)** Temporal dynamics of ZEB1/2 mRNA levels in a cell starting in an epithelial phenotype when exposed to a high level of an external EMT signal (I_ext = 100,000 molecules) (orange-shaded region) for WT1 coupled with the ELF3 network (pink curve), WT1 coupled with the core EMT circuit (no ELF3) (yellow curve), EMT–ELF3 circuit (blue curve) and core EMT circuit (no ELF3; no WT1: green curve). **D)** Phase diagrams for WT1 coupled with an ELF3 network driven by an external signal (l_ext) for varying strength of activation from WT1 to SNAIL. **E)** Same as D, but for varying threshold levels along of WT1 to activate SNAIL. **F)** Same as D, but for varying strength of inhibition of ZEB by ELF3. In D-F, different coloured regions show varied phases (combination of co-existing phenotypes).

We next mapped the temporal responses of these four circuits for a fixed value of I_ext signal. Among these four circuits, we found steady-state values of ZEB1/2 mRNA levels to be at a minimum in the presence of ELF3 and absence of WT1, and to be at a maximum in the presence of WT1 and absence of ELF3 (**Fig 5C**), thus supporting the ability of ELF3 to inhibit WT1-driven EMT. We next asked how specific interactions influence the ability of ELF3 to impact EMT dynamics. Increasing the strength of WT1-induced SNAIL activation – by either increasing the corresponding fold-change parameter (**Fig 5D**) or by reducing the threshold levels of WT1 needed to activate SNAIL (**Fig 5E**) – the region corresponding to a mesenchymal phenotype {M} expanded while that corresponding to an epithelial phenotype {E} decreased. These trends indicate that a stronger activation of SNAIL by WT1 can counteract the role of ELF3 as an EMT inhibitor. Conversely, an increase in the strength of ELF3-mediated ZEB1/2 inhibition leads to an expansion of the {E} region (only epithelial phenotype) accompanied by a shrinking of the {M} (only mesenchymal) and {H} (only hybrid E/M) regions (**Fig 5F**). Thus, ELF3 and WT1 can have opposite roles in enabling EMT progression.

Given the mutually-antagonistic relationship between WT1 and ELF3 in mediating EMT, we asked whether these factors demonstrated inverse trends in clinical data and correlated with patient outcomes. In breast cancer data sets, high WT1 levels correlated with improved relapse-free survival and overall survival (**Fig S6A-B**, GSE9893). However, this trend was reversed in other cancer types in which high WT1 associated with worse patient outcomes - colorectal cancer (relapse free survival: **Fig S6C-D**; GSE17536, GSE14333), lung cancer (overall survival: **Fig S6E-F**, GSE50081, GSE3141; relapse free survival: **Fig S6G**, GSE31210), ovarian cancer (overall survival: **Fig S6H**, GSE73614) and pancreatic cancer (overall survival: Fig **S6I**, TCGA-PAAD). Thus, in breast cancer, higher ELF3 or lower WT1 levels associated with worse outcomes, while in colorectal and ovarian cancer, lower ELF3 or higher WT1 levels had worse prognosis, reminiscent of the antagonistic role of ELF3 and WT1 in mediating phenotypic plasticity.

## Discussion

We propose ELF3 as a putative MET-TF, based on transcriptomic data analysis showcasing a strong association of ELF3 with an epithelial phenotype and its reversible reduction during EMT, as well as predictions from mechanism-based mathematical model for a network containing many core EMT/MET factors. These observations are in concordance with experimental data showing that silencing of ELF3 in NMuMG cells led to retention of a mesenchymal phenotype even when TGFβ was withdrawn, resulting in impaired MET (Sengez *et al*., 2019). Similarly, knockdown of ELF3 in biliary tract cancer cells resulted in upregulation of mesenchymal markers such as ZEB1/2, VIM and TWIST1 accompanied by the downregulation of KRT19 (Suzuki *et al*., 2021). Conversely, ELF3 over-expression in SKOV3 cells led to an inhibition of EMT (Yeung *et al*., 2017). Further, in gastric cancer, an antagonistic relation between ZEB1 and ELF3 was observed through their downstream targets, such as IRF6 (Li *et al*., 2019). In circulating tumor cells and in patient tumor biopsies, too (Liu *et al*., 2019; Balcik-Ercin *et al*., 2021), expression levels of ELF3 and ZEB1 were anti-correlated. Thus, similar to ELF5 (Chakrabarti *et al*., 2012; Wu *et al*., 2015; Yao *et al*., 2015), ELF3 may serve as an epithelial gatekeeper.

Besides being a potential epithelial gatekeeper, ELF3 is also involved in maintaining cancer cell stemness. In high-grade serous ovarian cancer (HGSOC), ELF3 forms a positive feedback loop with LGR4, which is involved in stem-cell renewal. Knockdown of ELF3 reduced tumorsphere formation (Wang *et al*., 2020). However, in bladder urothelial carcinoma, overexpressing ELF3 repressed tumor-sphere formation despite antagonizing EMT (Na *et al*., 2022). Hence, the interplay between ELF3, EMT and stemness appears to be context-specific, reminiscent of recent observations associating various stages of EMT with enhanced tumor-initiation potential in many cancer types (Pastushenko *et al*., 2018; Kröger *et al*., 2019; Pasani *et al*., 2021; Brown *et al*., 2022). Such context-specific associations may underlie lineage-restricted roles of ELF3 as a tumor suppressor or an oncogene, depending on cancer cell lineage and/or differentiation status (Enfield *et al*., 2019).

In addition to stemness, the role of ELF3 in conferring therapy resistance has been investigated. ELF3 has been reported to be upregulated in NSCLC cells resistant to <u>the</u> PARP inhibitor, olaparib (Wang *et al*., 2021). Further, in NSCLC cells, treatment with auranofin reduced ELF3 levels and induced cell death (Lee *et al*., 2021). Similarly, ELF3 overexpression in ovarian cancer cells reduced their sensitivity to cisplatin (Liu *et al*., 2021). Future investigations should interrogate the coupled dynamics of ELF3, EMT and resistance to specific therapies, similar to our observations that ELF3 is associated with an epithelial and tamoxifen-resistant cell-state.

Our analysis also revealed that while ELF3 may show stronger association with an epithelial state when compared with KLF4, but its effects in inducing MET were found to be weaker than GRHL2. GRHL2 is a pioneering transcription factor that can bind to closed chromatin and initiate its opening (Chen *et al*., 2018; Balsalobre and Drouin, 2022). Although our mechanism-based mathematical model does not incorporate epigenetic interactions, the ability of GRHL2 to influence chromatin-level reprogramming further elevates its potency as a strong MET inducer (Chung *et al*., 2019). GRHL2 has also been reported to be lineage-specific driver of reprogrammed estrogen signaling and an enabler of endocrine resistance in ER+ breast cancer (Cocce *et al*., 2019; Kumegawa *et al*., 2022). While GRHL2 overexpression was sufficient to induce MET in mesenchymal MDA-MB-231 breast cancer cells, it failed to do so in RD sarcoma cells (Somarelli *et al*., 2016). Further analysis of EMT/MET inducing transcription factors should thus consider tissue lineage as a crucial axis, because of varying potency of these factors in facilitating lineage-restricted phenotypic plasticity.

## Materials and Methods

### Mathematical modeling

A system of coupled ordinary differential equations were employed to understand the dynamics of the ELF3 coupled EMT circuit comprising of miR-200, SNAIL, ZEB, SLUG and KLF4 (**Fig 2E**). The following generic chemical rate equation describes the level of a protein, mRNA or micro-RNA (X):

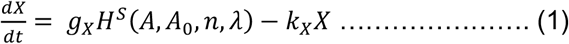

where g_X_ represents the basal rate of production, transcriptional/translational/post-translational regulations is represented by the terms multiplied by g_X._ – one or more shifted Hills function (*H*^*S*^(*A, A*_*0*_, *n, λ*)) that describe the interactions among the species in the system. The degradation of species (X) is assumed to follow first-order kinetics and thus defined by the term k_X_X.

The complete set of equations and parameters are presented in the Supplementary Material. Bifurcation diagrams were drawn in MATLAB (MathWorks Inc.) using the continuation software package MATCONT (Dhooge *et al*., 2008).

### RACIPE (random network simulation)

Random Circuit Perturbation (RACIPE) is a simulation framework that extensively explore the possible multistable properties of a given gene regulatory network (Huang *et al*., 2017). Based on the gene regulatory network topology, x coupled ordinary differential equations (ODEs) are simulated to obtain the multistable properties of the gene regulatory network (x is the number of nodes/genes in the network). The parameters for the set of coupled ordinary differential equations are sampled randomly from pre-defined ranges that ensures a robust sampling of a large parameter space that can represent the overall dynamical properties of the gene regulatory network. The program samples 10000 sets of parameters and for each parameter set, RACIPE initialises the system with a random set of initial conditions (n = 100) for each node in the network. The parameterised set of ODEs are then solved using the Eulers method to obtain one or many steady states that represent the attractors that are enabled by each parameter set. The steady state expression values are then z-normalised for principal component analysis (PCA) and hierarchical clustering analysis. The perturbation analysis was done by performing RACIPE analysis on a gene regulatory network by either over expressing (OE) or down expressing (DE) a specified node by x-fold (i.e. the production rate of that particular gene is increase by x-folds and the steady state values are computed for the set of coupled ODEs). The Z-score normalisation of these perturbation data was done with respect to the control case where none of the production rates were altered. The proportion of phenotypes in each case were then computed over three replicates of in-silico perturbations to assess for statistical significance.

### Gene expression datasets

Gene expression datasets were downloaded using the GEOquery R Bioconductor package (Davis and Meltzer, 2007). The datasets were pre-processed for each sample and gene-wise expression data was obtained from probe-wise expression matrix using R (version 4.0.0). To calculate the Epithelial and/or the mesenchymal scores for bulk RNA seq data, we used the ssGSEA functionality to estimate the activity of either the epithelial and/or the mesenchymal set of genes for each sample in the corresponding datasets. The epithelial and mesenchymal gene lists were obtained from (Tan *et al*., 2014). The Hallmark EMT gene set was obtained from MSigDB (Liberzon *et al*., 2011). For the single cell RNA seq dataset, GSE147405 (Cook and Vanderhyden, 2020), imputation of gene expression values was performed using MAGIC (van Dijk *et al*., 2018) before plotting the expression levels of ELF3, KLF4 and GRHL2. Imputed values were also used to calculate the activity of the gene signatures such as the Hallmark EMT signature using AUCell (Aibar *et al*., 2017). We computed the Spearman correlation coefficients and used corresponding p-values to gauge the strength of correlations for all correlation analysis. For statistical comparison between discrete groups, we used a two-tailed Student’s t-test under the assumption of unequal variances and computed significance.

### Kaplan-Meier analysis

Kaplan-Meier analysis for respective datasets was performed using ProgGene (Goswami and Nakshatri, 2014). The number of samples showing high and low expression levels of ELF3 and WT1 are indicated in the SI table.

## Author Contributions

MKJ conceptualized research; MKJ and JAS supervised research; ARS, SS, PM, SG, VAK, YM, SR and IM performed research and analyzed data. All authors contributed to writing and review of the manuscript.

## Conflict of Interest

The authors declare no conflict of interest.

## Funding

This work was supported by Ramanujan Fellowship (SB/S2/RJN-049/2018) awarded to MKJ by Science and Engineering Research Board (SERB), Department of Science and Technology (DST), Government of India. JAS is supported by the Department of Defense (W81XWH-18-1-0189) and NCI 1R01CA233585-03. SS was awarded Prime Ministers’ Research Fellowship (PMRF) by Government of India.

## Notes

### Competing Interest Statement

The authors have declared no competing interest.

