## Supplementary Figures & Text for "The ELF3 transcription factor is associated with an epithelial phenotype and represses epithelial-mesenchymal transition"

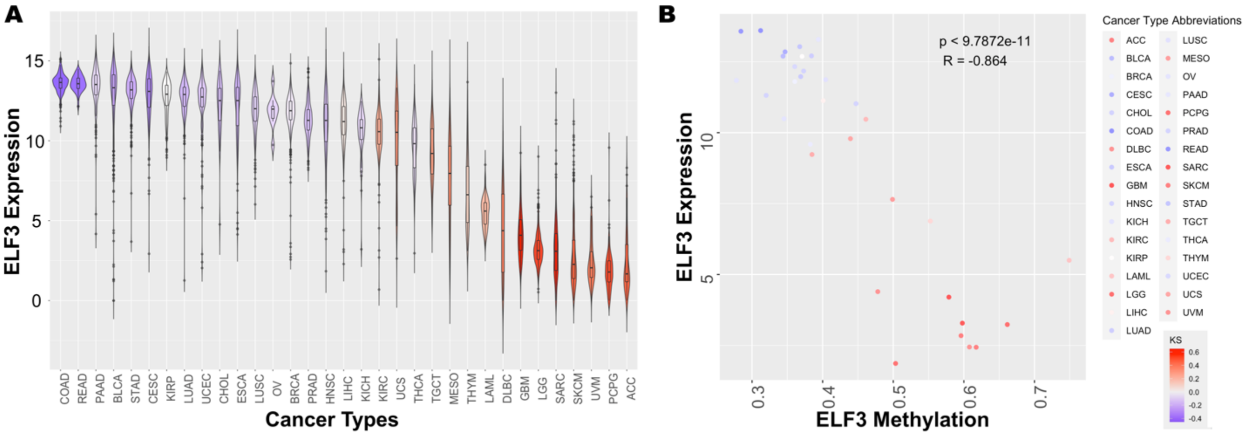


***Fig S1: TCGA data for ELF3. A)*** *ELF3 expression levels in TCGA cancer samples ordered by low KS score to high KS score.* ***B)*** *Scatter plot for ELF3 expression and its methylation status in TCGA cancer types. Each dot (cancer type) colored by KS score.*

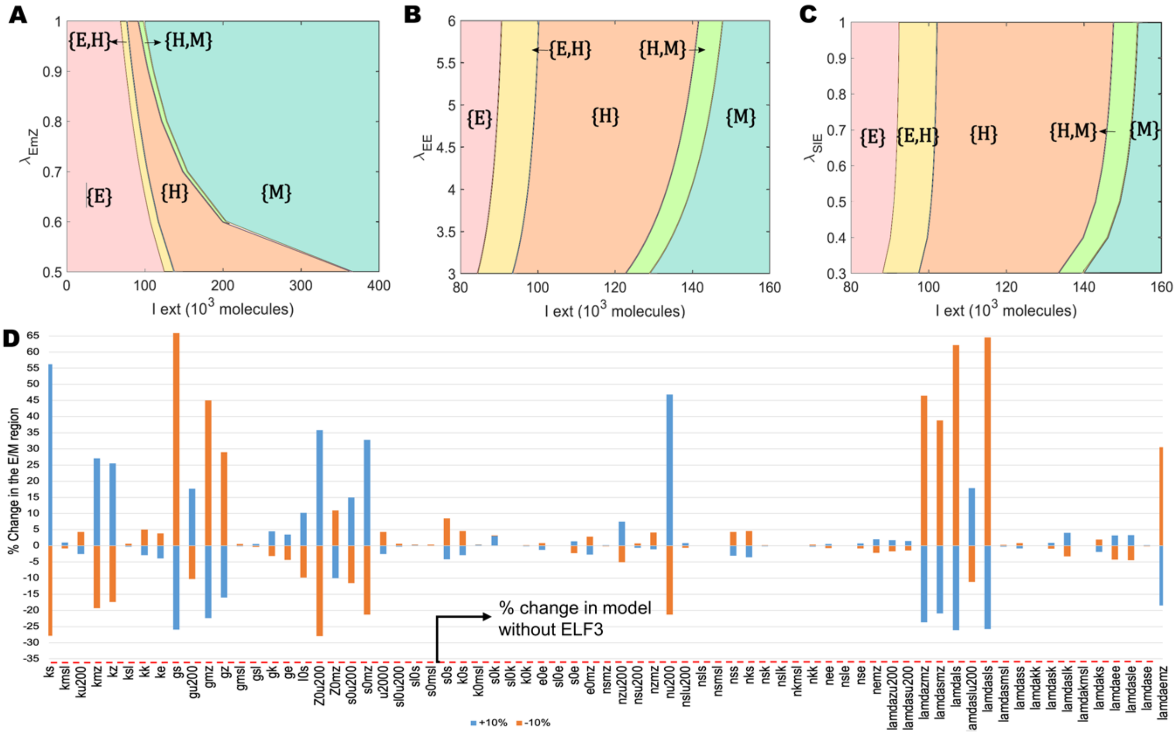


***Fig S2: ELF3 delays the onset of EMT. (A)*** *Phase diagrams for the ELF3 network driven by an external signal (I_ext) for varying strength of interactions along the ELF3-ZEB axis* ***(B)*** *Phase diagrams for the ELF3 network driven by an external signal (I_ext) for varying strength of ELF3 self-activation.* ***(C)*** *Phase diagrams for the ELF3 network driven by an external signal (I_ext) for varying strength of interactions along the Slug-ELF3 axis (D) Sensitivity analysis of parameters for the ELF3 coupled EMT circuit (Fig 2E) indicating percent change in the I_ext interval for which the hybrid E/M state exists. The red dotted line indicates the percent change in the stable hybrid region in the absence of ELF3 (core network) when compared to the coupled network.*

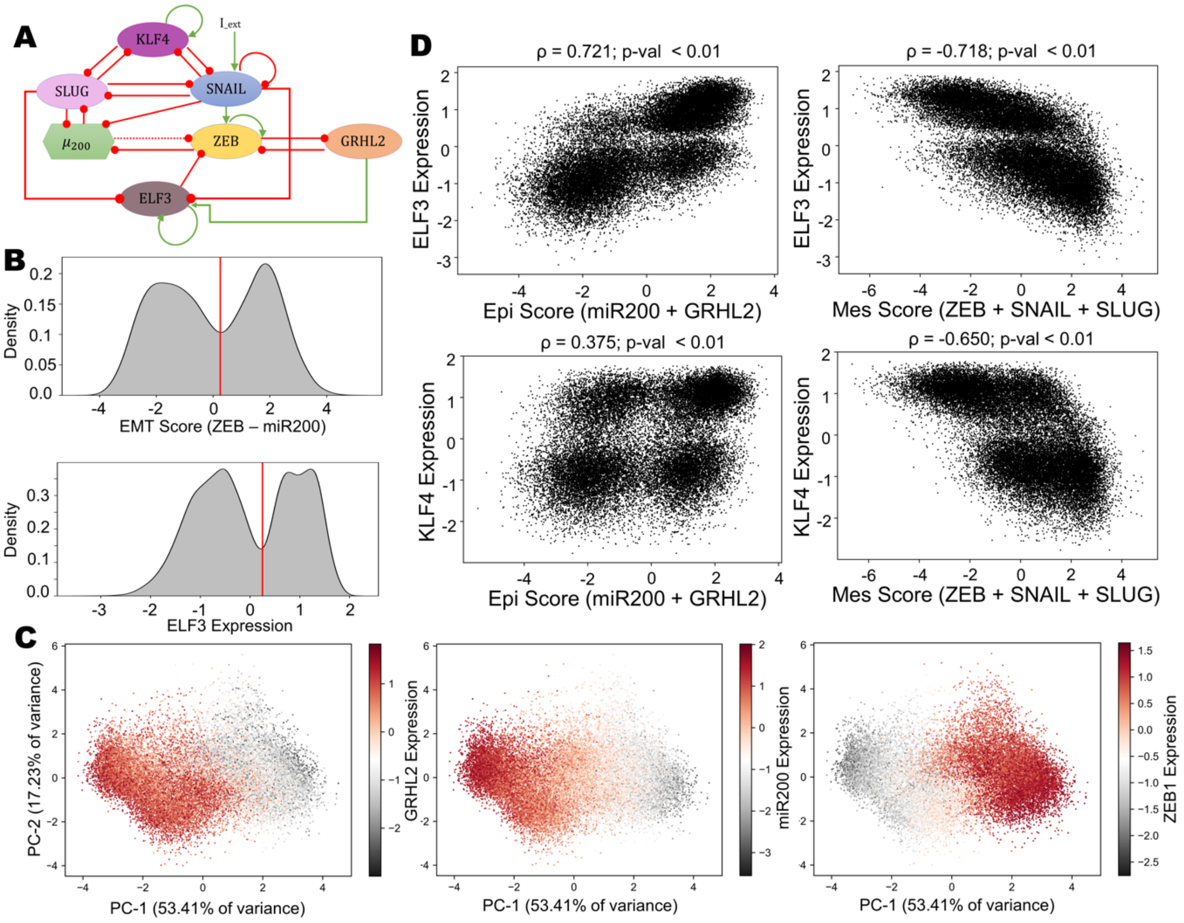


***Fig S3: ELF3 is a stronger inducer of MET than KLF4. (A)*** *Gene regulatory network showing the regulation between epithelial and mesenchymal genes. Green arrows indicate activatory links and Red hammers indicate inhibitory links.* ***(B)*** *Kernel density estimate plots of EMT score (ZEB – miR200) (top panel) and ELF3 z-normalized expression (bottom panel). The red vertical line shows the approximate position of the minima of the largely bimodal distributions. PCA scatter plot of all steady states of RACIPE colored by (i) the EMT score defined as ZEB – miR200, (ii) SLUG expression (iii) ELF3 expression and (iv) KLF4 expression.* ***(C)*** *Scatterplot of Epithelial scores (GRHL2 + miR200) and Mesenchymal scores (ZEB + SNAIL +SLUG) with ELF3 and ELF4 levels of steady state solutions from RACIPE. The spearman correlation coefficient and the corresponding p-values have been mentioned.* ***(D)*** *PCA scatter plot of all steady states of RACIPE colored by GRHL2 Expression (leftmost panel), miR200 expression (center panel) and ZEB expression (rightmost panel).*

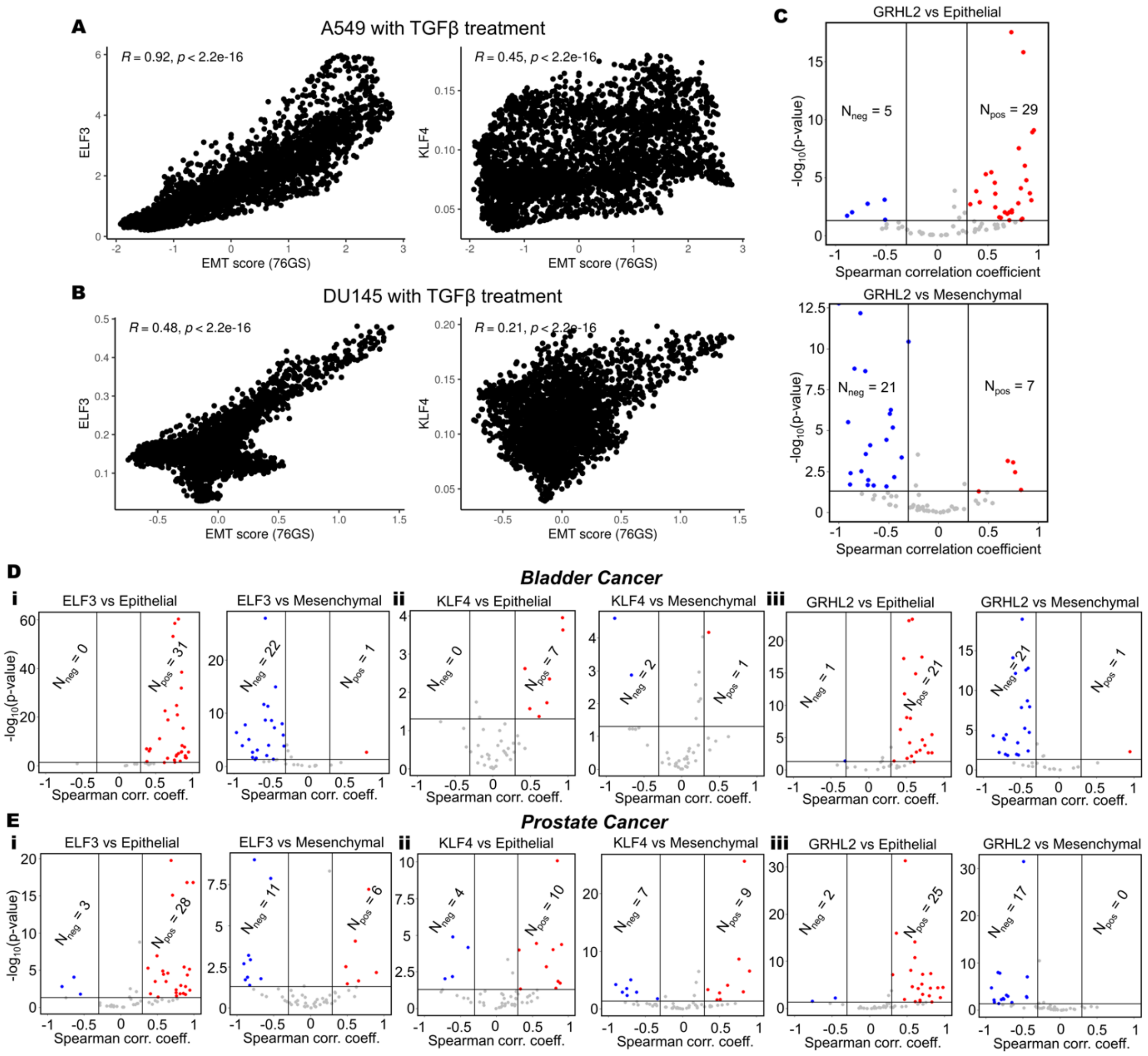


***Fig S4: ELF3 shows stronger trends as compared to KLF4 with an epithelial behavior**. Scatter plots showing correlations of imputed ELF3 and KLF4 expression with 76GS EMT score (the higher the 76GS score, the more epithelial the sample) in two cancer cell lines **A)** A549 (lung cancer) and **B)** DU145 (prostate cancer) when treated with TGFβ (GSE147405). Spearman correlation coefficient and the corresponding p-values have been mentioned. Imputed gene expression values were calculated using the MAGIC algorithm for different cell lines separately. **C)** Volcano plots showing correlation of GRHL2 expression levels with ssGSEA epithelial and mesenchymal scores in a meta-analysis of breast cancer datasets. Each dot represents a dataset. R < - 0.3, p< 0.05 or R > 0.3, p < 0.05 are counted as statistically significant cases. $N_{neg}$ denotes number of datasets for which a negative correlation (blue dots) is observed, $N_{pos}$ denotes number of datasets for which a positive correlation (red dots) is observed between the two corresponding expression levels or ssGSEA scores. **D)** Same as C) but for bladder cancer. **E)** Same as C) but for prostate cancer. Panels D and E show results for KLF4, ELF3 and GRHL2.*

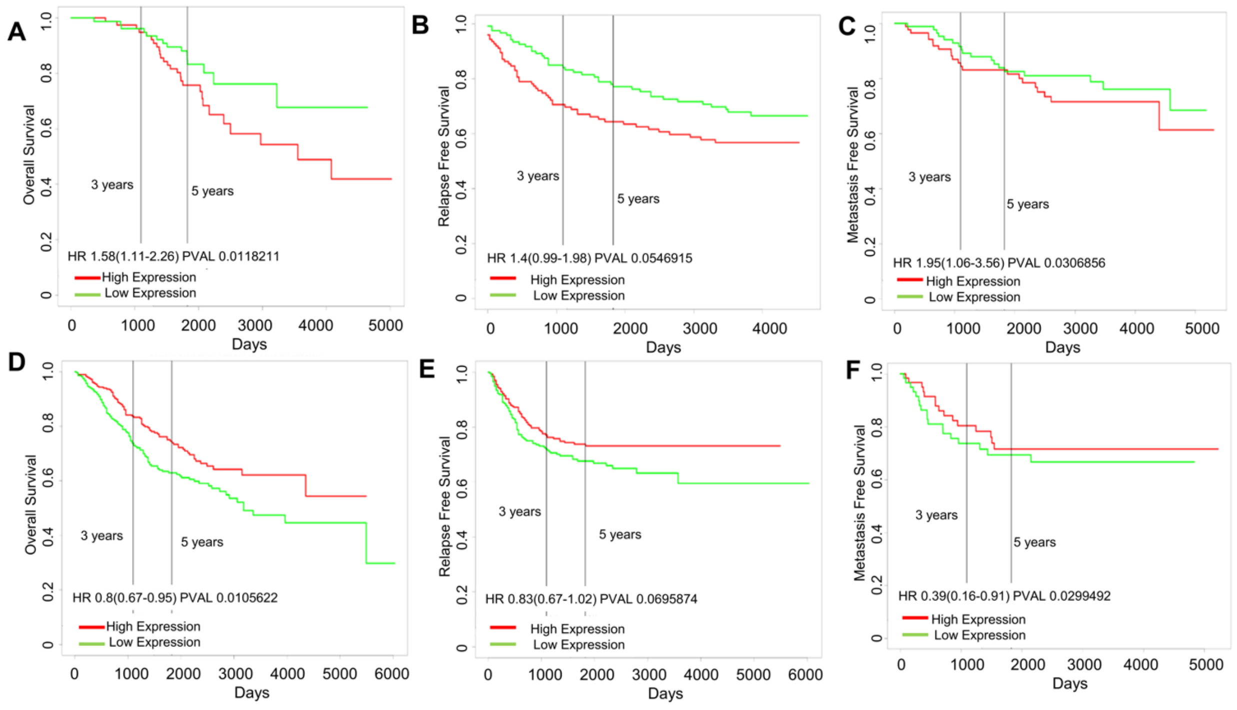


***Fig S5: ELF3 correlates with patient survival in a cancer-specific manner. (A)*** *Trends in breast cancer samples. (i) overall survival (GSE9893) (ii) relapse-free survival (GSE4922) (iii) metastasis-free survival (GSE6532)* ***(B)*** *Trends in colorectal cancer samples. (i) overall survival (GSE39582) (ii) relapse-free survival (GSE395824) (iii) metastasis-free survival (GSE28722).*

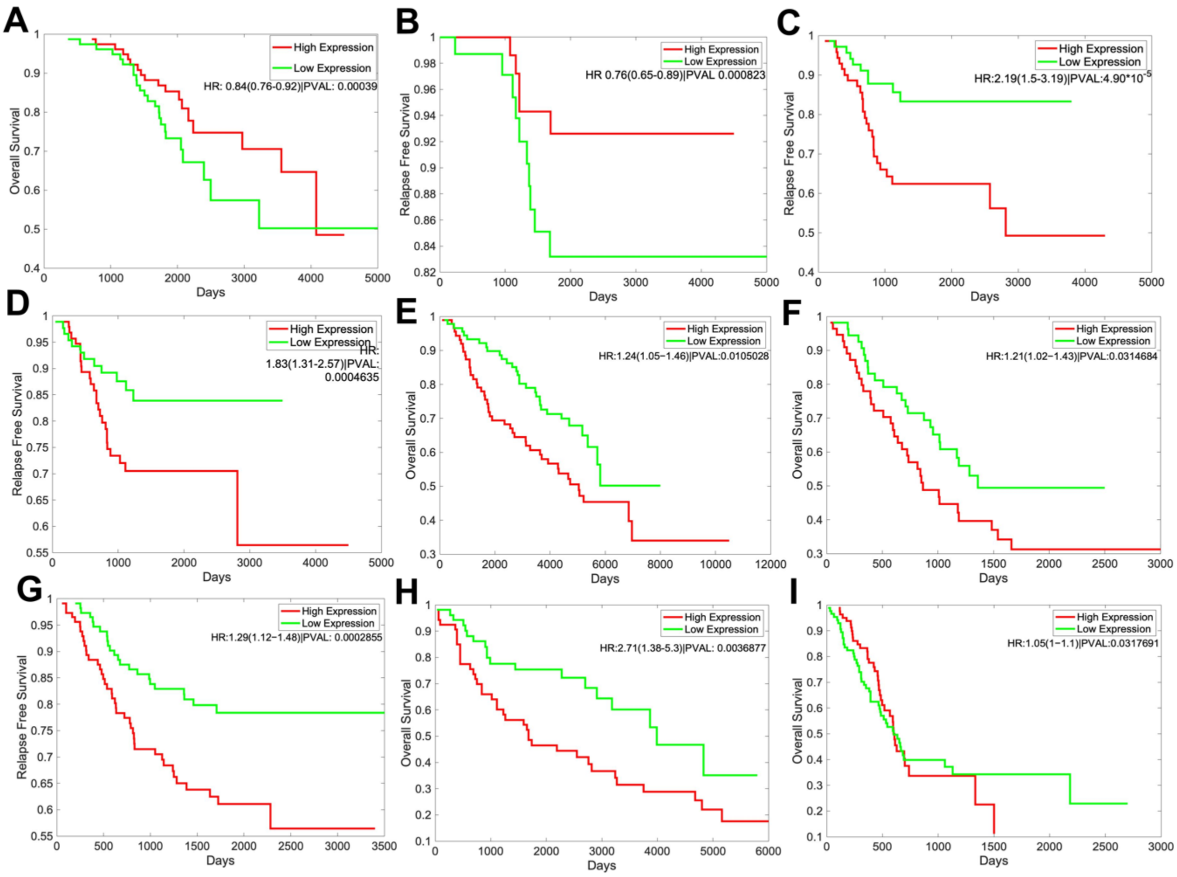


***Fig S6: WT1 correlates with patient survival in a cancer-specific manner.*** *(A) Overall survival in Breast cancer sample (GSE9893). (B) Relapse free survival in breast cancer sample (GSE9893). (C) (D) Relapse free survival in colorectal cancer samples (GSE17536, GSE14333) (E) (F) Overall survival in lung cancer samples (GSE50081, GSE314) (G) Relapse free survival in lung cancer sample (GSE31210) (H) Overall survival in ovarian cancer sample (GSE73614) (I) Overall survival in pancreatic cancer sample (TCGA-PAAD)*

**Supplementary Information**

Coupled ordinary differential equations were used to describe the dynamics of the molecular species present in the EMT regulatory circuits shown in Fig 2E and 5A.

Equations describing the dynamics of the core circuit comprising of ZEB, SNAIL, SLUG, miR200 and KLF4

$$\frac{d\mu_{200}}{dt} = g_{\mu_{200}} H^s(Z,\lambda_{Z,\mu_{200}}) H^s(S,\lambda_{S,\mu_{200}}) H^s(Sl,\lambda_{Sl,\mu_{200}}) - m_Z Y_\mu(\mu_{200}) - m_{Sl} Y_\mu(\mu_{200}) - k_{\mu_{200}}\mu_{200}$$

$$\frac{dm_Z}{dt} = g_{m_Z} H^s(Z,\lambda_{Z,m_Z}) H^s(S,\lambda_{S,m_Z}) - m_Z Y_Z(\mu_{200}) - k_{m_Z} m_Z$$

$$\frac{dZ}{dt} = g_Z m_Z L(\mu_{200}) - k_Z Z$$

$$\frac{dS}{dt} = g_S H^s(I,\lambda_{I,S}) H^s(Sl,\lambda_{Sl,S}) H^s(S,\lambda_{S,S}) H^s(K,\lambda_{K,S}) - k_S S$$

$$\frac{dm_{Sl}}{dt} = g_{m_{Sl}} H^s(S,\lambda_{S,m_{Sl}}) H^s(K,\lambda_{K,m_{Sl}}) - m_{Sl} Y_Z(\mu_{200}) - k_{m_{Sl}} m_{Sl}$$

$$\frac{dSl}{dt} = g_{Sl} m_{Sl} L(\mu_{200}) - k_{Sl} Sl$$
$$\frac{dK}{dt} = g_K H^s(\mathrm{K}, \lambda_{K,K}) H^s(Sl, \lambda_{Sl,K}) H^s(S, \lambda_{S,K}) - k_K K$$

Equations describing the dynamics of the ELF3 circuit comprising of ZEB, SNAIL, SLUG, miR200, KLF4 and ELF3

$$\frac{d\mu_{200}}{dt} = g_{\mu_{200}} H^s(\mathrm{Z}, \lambda_{Z,\mu_{200}}) H^s(S, \lambda_{S,\mu_{200}}) H^s(Sl, \lambda_{Sl,\mu_{200}}) - m_Z Y_\mu(\mu_{200}) - m_{Sl} Y_\mu(\mu_{200}) - k_{\mu_{200}} \mu_{200}$$
$$\frac{dm_Z}{dt} = g_{m_Z} H^s(\mathrm{Z}, \lambda_{Z,m_Z}) H^s(S, \lambda_{S,m_Z}) H^s(\mathrm{E}, \lambda_{E,m_Z}) - m_Z Y_Z(\mu_{200}) - k_{m_Z} m_Z$$
$$\frac{dZ}{dt} = g_Z m_Z L(\mu_{200}) - k_Z Z$$
$$\frac{dS}{dt} = g_S H^s(\mathrm{I}, \lambda_{I,S}) H^s(Sl, \lambda_{Sl,S}) H^s(S, \lambda_{S,S}) H^s(K, \lambda_{K,S}) - k_S S$$
$$\frac{dm_{Sl}}{dt} = g_{m_{Sl}} H^s(S, \lambda_{S,m_{Sl}}) H^s(K, \lambda_{K,m_{Sl}}) - m_{Sl} Y_Z(\mu_{200}) - k_{m_{Sl}} m_{Sl}$$
$$\frac{dSl}{dt} = g_{Sl} m_{Sl} L(\mu_{200}) - k_{Sl} Sl$$
$$\frac{dK}{dt} = g_K H^s(\mathrm{K}, \lambda_{K,K}) H^s(Sl, \lambda_{Sl,K}) H^s(S, \lambda_{S,K}) - k_K K$$
$$\frac{dE}{dt} = g_E H^s(\mathrm{E}, \lambda_{E,E}) H^s(S, \lambda_{S,E}) H^s(Sl, \lambda_{Sl,E}) - k_E E$$

Equations describing the dynamics of the ELF3+ERα circuit comprising of ZEB, SNAIL, SLUG, miR200, KLF4, ELF3, ERα-66 and ERα36

$$\frac{d\mu_{200}}{dt} = g_{\mu_{200}} H^s(\mathrm{Z}, \lambda_{Z,\mu_{200}}) H^s(S, \lambda_{S,\mu_{200}}) H^s(Sl, \lambda_{Sl,\mu_{200}}) - m_Z Y_\mu(\mu_{200}) - m_{Sl} Y_\mu(\mu_{200}) - k_{\mu_{200}} \mu_{200}$$
$$\frac{dm_Z}{dt} = g_{m_Z} H^s(\mathrm{Z}, \lambda_{Z,m_Z}) H^s(S, \lambda_{S,m_Z}) H^s(\mathrm{E}, \lambda_{E,m_Z}) H^s(ER_{36}, \lambda_{ER_{36},m_Z}) - m_Z Y_Z(\mu_{200}) - k_{m_Z} m_Z$$
$$\frac{dZ}{dt} = g_Z m_Z L(\mu_{200}) - k_Z Z$$
$$\frac{dS}{dt} = g_S H^s(\mathrm{I}, \lambda_{I,S}) H^s(Sl, \lambda_{Sl,S}) H^s(S, \lambda_{S,S}) H^s(K, \lambda_{K,S}) - k_S S$$
$$\frac{dm_{Sl}}{dt} = g_{m_{Sl}} H^s(S, \lambda_{S,m_{Sl}}) H^s(ER_{66}, \lambda_{ER_{66},m_{Sl}}) H^s(K, \lambda_{K,m_{Sl}}) - m_{Sl} Y_Z(\mu_{200}) - k_{m_{Sl}} m_{Sl}$$
$$\frac{dSl}{dt} = g_{Sl} m_{Sl} L(\mu_{200}) - k_{Sl} Sl$$
$$\frac{dK}{dt} = g_K H^s(\mathrm{K}, \lambda_{K,K}) H^s(Sl, \lambda_{Sl,K}) H^s(S, \lambda_{S,K}) - k_K K$$
$$\frac{dE}{dt} = g_E H^s(\mathrm{E}, \lambda_{E,E}) H^s(S, \lambda_{S,E}) H^s(Sl, \lambda_{Sl,E}) H^s(ER_{66}, \lambda_{ER_{66},E}) - k_E E$$
$$\frac{dER_{66}}{dt} = g_{ER_{66}} H^s(ER_{66}, \lambda_{ER_{66},ER_{66}}) H^s(\mathrm{E}, \lambda_{E,ER_{66}}) H^s(Z, \lambda_{Z,ER_{66}}) - k_{ER_{66}} ER_{66}$$
$$\frac{dER_{36}}{dt} = g_{ER_{36}} H^s(ER_{66}, \lambda_{ER_{66},ER_{36}}) - k_{ER_{36}} ER_{36}$$

Equations describing the dynamics of the WT1 circuit comprising of ZEB, SNAIL, SLUG, miR200, KLF4 and WT1

$$\frac{d\mu_{200}}{dt} = g_{\mu_{200}} H^s(\mathrm{Z}, \lambda_{Z,\mu_{200}}) H^s(S, \lambda_{S,\mu_{200}}) H^s(Sl, \lambda_{Sl,\mu_{200}}) - m_Z Y_\mu(\mu_{200}) - m_{Sl} Y_\mu(\mu_{200}) - k_{\mu_{200}} \mu_{200}$$
$$\frac{dm_Z}{dt} = g_{m_Z} H^s(\mathrm{Z}, \lambda_{Z,m_Z}) H^s(S, \lambda_{S,m_Z}) - m_Z Y_Z(\mu_{200}) - k_{m_Z} m_Z$$
$$\frac{dZ}{dt} = g_Z m_Z L(\mu_{200}) - k_Z Z$$
$$\frac{dS}{dt} = g_S H^s(\mathrm{I}, \lambda_{I,S}) H^s(Sl, \lambda_{Sl,S}) H^s(S, \lambda_{S,S}) H^s(K, \lambda_{K,S}) H^s(\mathrm{W}, \lambda_{W,S}) - k_S S$$
$$\frac{dm_{Sl}}{dt} = g_{m_{Sl}} H^s(S, \lambda_{S,m_{Sl}}) H^s(K, \lambda_{K,m_{Sl}}) H^s(\mathrm{W}, \lambda_{W,m_{Sl}}) - m_{Sl} Y_Z(\mu_{200}) - k_{m_{Sl}} m_{Sl}$$
$$\frac{dSl}{dt} = g_{Sl} m_{Sl} L(\mu_{200}) - k_{Sl} Sl$$
$$\frac{dK}{dt} = g_K H^s(\mathrm{K}, \lambda_{K,K}) H^s(Sl, \lambda_{Sl,K}) H^s(S, \lambda_{S,K}) - k_K K$$
$$\frac{dW}{dt} = g_W H^s(\mathrm{W}, \lambda_{W,W}) - k_W W$$

Equations describing the dynamics of the (ELF3+WT1) circuit comprising of ZEB, SNAIL, SLUG, miR200, KLF4, ELF3 and WT1

$$\frac{d\mu_{200}}{dt} = g_{\mu_{200}} H^s(\mathrm{Z}, \lambda_{Z,\mu_{200}}) H^s(S, \lambda_{S,\mu_{200}}) H^s(Sl, \lambda_{Sl,\mu_{200}}) - m_Z Y_\mu(\mu_{200}) - m_{Sl} Y_\mu(\mu_{200}) - k_{\mu_{200}} \mu_{200}$$
$$\frac{dm_Z}{dt} = g_{m_Z} H^s(\mathrm{Z}, \lambda_{Z,m_Z}) H^s(S, \lambda_{S,m_Z}) H^s(\mathrm{E}, \lambda_{E,m_Z}) - m_Z Y_Z(\mu_{200}) - k_{m_Z} m_Z$$
$$\frac{dZ}{dt} = g_Z m_Z L(\mu_{200}) - k_Z Z$$
$$\frac{dS}{dt} = g_S H^s(\mathrm{I}, \lambda_{I,S}) H^s(Sl, \lambda_{Sl,S}) H^s(S, \lambda_{S,S}) H^s(K, \lambda_{K,S}) H^s(\mathrm{W}, \lambda_{W,S}) - k_S S$$
$$\frac{dm_{Sl}}{dt} = g_{m_{Sl}} H^s(S, \lambda_{S,m_{Sl}}) H^s(K, \lambda_{K,m_{Sl}}) H^s(\mathrm{W}, \lambda_{W,m_{Sl}}) - m_{Sl} Y_Z(\mu_{200}) - k_{m_{Sl}} m_{Sl}$$
$$\frac{dSl}{dt} = g_{Sl} m_{Sl} L(\mu_{200}) - k_{Sl} Sl$$
$$\frac{dK}{dt} = g_K H^s(\mathrm{K}, \lambda_{K,K}) H^s(Sl, \lambda_{Sl,K}) H^s(S, \lambda_{S,K}) - k_K K$$
$$\frac{dE}{dt} = g_E H^s(\mathrm{E}, \lambda_{E,E}) H^s(S, \lambda_{S,K}) H^s(Sl, \lambda_{Sl,K}) - k_E E$$

$$\frac{dW}{dt} = g_W H^s(\mathrm{W}, \lambda_{W,W}) - k_W W$$

where $g_X$ is the corresponding innate production rate and $k_X$ is the innate degradation rate. $m_Z L(\mu_{200})$ is the net translation rate, $m_Z Y_m(\mu_{200})$ is the total ZEB mRNA active degradation rate and $m_Z Y_\mu(\mu_{200})$ is the total miR active degradation rate. $H^S$ is the shifted Hill function, defined as
$H^S(B,\lambda) = H^-(B)+\lambda H^+(B)$ ,
$H^-(B) = 1/ [1+(B / B_0)^{nB}]$ ,
$H^+(B) = 1-H^-(B)$,
λ is the fold change from the basal synthesis rate due to protein B. λ >1 for activators, while λ<1 for inhibitors.

λ<1 for inhibitors.

**Parameter Estimation:**
The model parameters were adopted from previously published literature for the molecular species of the core circuit (I, miR-200, Snail, Zeb, Slug) and KLF4 interactions, as given below:

| Parameter | Value | Reference |
| --- | --- | --- |
| ks | 0.125 | (Lu et al., 2013) |
| ku200 | 0.05 | (Lu et al., 2013) |
| kmz | 0.5 | (Lu et al., 2013) |
| kz | 0.1 | (Lu et al., 2013) |
| kmsl | 0.5 | Estimated |
| ksl | 0.1155 | (Lu et al., 2013) |
| kk | 0.1732 | (Lu et al., 2013) |
| ke | 0.125 | Estimated |
| kw | 0.2722 | (Scharnhorst et al., 1999) |
| gs | 18000 | (Lu et al., 2013) |
| gu200 | 2100 | (Lu et al., 2013) |
| gmz | 11 | (Lu et al., 2013) |
| gz | 100 | (Lu et al., 2013) |
| gmsl | 90 | Estimated |
| gsl | 50000 | Estimated |
| gk | 50000 | Estimated |
| ge | 50000 | Estimated |
| gw | 10000 | Estimated |
| I0s | 100000 | (Jolly et al., 2017) |
| Z0u200 | 220000 | (Lu et al., 2013) |
| Z0mz | 27500 | (Lu et al., 2013) |
| s0u200 | 180000 | (Lu et al., 2013) |
| s0mz | 180000 | (Lu et al., 2013) |
| u2000 | 10000 | (Lu et al., 2013) |
| sl0u200 | 220000 | Estimated |
| sl0msl | 150000 | Estimated |
| sl0s | 225000 | Estimated |
| s0msl | 180000 | Estimated |
| s0s | 300000 | Estimated |
| k0s | 275000 | Estimated |
| k0msl | 300000 | Estimated |
| s0k | 180000 | Estimated |
| sl0k | 250000 | Estimated |
| k0k | 275000 | Estimated |
| e0e | 200000 | Estimated |
| sl0e | 220000 | Estimated |
| s0e | 250000 | Estimated |
| e0mz | 180000 | Estimated |
| w0s | 350000 | Estimated |
| w0msl | 220000 | Estimated |
| w0w | 280000 | Estimated |
| nsmz | 2 | (Lu et al., 2013) |
| nIs | 2 | (Jolly et al., 2017) |

| | | |
|---|---|---|
| nzu200 | 3 | (Lu et al., 2013) |
| nsu200 | 2 | (Lu et al., 2013) |
| nzmz | 2 | (Lu et al., 2013) |
| nu200 | 6 | (Lu et al., 2013) |
| nslu200 | 1 | (Y. N. Liu et al., 2013) |
| nsls | 3 | (Chen and Gridley, 2013) |
| nsmsl | 1 | (Chen and Gridley, 2013) |
| nss | 5 | (Chen and Gridley, 2013) |
| nks | 2 | (Yori et al., 2011) |
| nsk | 2 | estimated |
| nslk | 4 | (Y.-N. Liu et al., 2012) |
| nkmsl | 2 | (Y.-N. Liu et al., 2012) |
| nkk | 3 | (Mahatan et al., 1999) |
| nee | 3 | (Kopp et al., 2007) |
| nsle | 4 | Estimated |
| nse | 4 | Estimated |
| nemz | 2 | (Suzuki et al., 2021) |
| nww | 8 | (Hewitt et al., 1996) |
| nwmsl | 1 | (Takeichi et al., 2013) |
| nws | 1 | (Martínez-Estrada et al., 2010) |
| lamdazu200 | 0.1 | (Lu et al., 2013) |
| lamdasu200 | 0.1 | (Lu et al., 2013) |
| lamdazmz | 7.5 | (Lu et al., 2013) |
| lamdasmz | 10 | (Lu et al., 2013) |
| lamdals | 3 | (Jolly et al., 2017) |
| lamdaslu200 | 0.4 | (Y. N. Liu et al., 2013) |
| lamdasls | 0.5 | (Nakamura et al., 2018) |
| lamdasmsl | 0.5 | (Nakamura et al., 2018) |
| lamdass | 0.4 | (Peiró et al., 2006) |
| lamdakk | 2 | (Dang, 2002) |
| lamdask | 0.25 | (Z. Li et al., 2018) |
| lamdaslk | 0.5 | (Y. N. Liu et al., 2013) |
| lamdakmsl | 0.25 | (Y. N. Liu et al., 2013) |
| lamdaks | 0.5 | (Yori et al., 2011) |
| lamdaee | 4 | (Q. Li et al., 2021) |
| lamdasle | 0.3 | (Lyons et al., 2008) |
| lamdase | 0.25 | (Lyons et al., 2008) |
| lamdaemz | 0.675 | (Suzuki et al., 2021) |
| lamdaws | 6 | (Sampson et al., 2014) |
| lamdawmsl | 0.5 | (Takeichi et al., 2013) |
| lamdaww | 0.4 | (Han et al., 2007) |

### Datasets Used in Kaplan-Meier analysis

| Datasets used for ELF3 analysis | | |
|---|---|---|
| **Dataset** | **n(High)** | **n(low)** |
| GSE3494 | 118 | 118 |
| GSE4922 | 124 | 124 |
| GSE48408 | 82 | 82 |
| GSE16125 | 16 | 16 |
| GSE28814 | 61 | 61 |
| GSE9893 | 77 | 77 |
| GSE6532 | 88 | 90 |
| GSE39582 | 288 | 284 |
| GSE28722 | 63 | 62 |
| **Datasets used for WT1 analysis** | | |
| **Dataset** | **n(High)** | **n(low)** |
| GSE9893 | 77 | 77 |
| GSE17536 | 71 | 69 |
| GSE14333 | 94 | 93 |
| GSE50081 | 91 | 90 |
| GSE3141 | 56 | 55 |
| GSE31210 | 113 | 113 |
| GSE73614 | 53 | 53 |
| TCGA-PAAD | 85 | 85 |
